# Mapping bioimage-analysis software adoption in microscopy

**DOI:** 10.64898/2026.10.21.760457

**Authors:** Tayla Shakespeare, Bruno M. Saraiva, Inês Cunha, Caron Jacobs, Guillaume Jacquemet, Ricardo Henriques

## Abstract

Most quantitative microscopy measurements pass through image-analysis software, and different programs can return different measurements from the same images. Which programs researchers rely on therefore matters for reproducibility; however, this information lies scattered across the methods sections of the microscopy literature. We tracked 32 software entries across 1.7 million full-text microscopy papers published between 2010 and 2025. In 2010, 15% of these papers mentioned at least one of the 23 analysis packages in our aggregate; by 2025, 31% did. Mentions stayed concentrated on a few general-purpose platforms, led by ImageJ/Fiji. Papers mentioning newer packages more often use the vocabulary of particular fields, such as pathology terms with QuPath and electron-microscopy terms with napari. Deep-learning methods are usually mentioned alongside an established platform; in sampled papers, StarDist often ran inside Fiji and Cellpose on its own. Many papers mention a package without citing it, so citation counts understate how often packages appear in published work; authors should make that use visible by citing each package and reporting how they ran it.

## Introduction

Modern microscopy produces multi-dimensional digital images, and extracting biological measurements from them relies on computation. In methods such as single-molecule localisation, structured illumination and fluctuation nanoscopy, computation is inseparable from image formation itself (1–4). As bioimage datasets grow, their analysis depends increasingly on scalable data access and interoperable formats (5, 6). Biological conclusions therefore depend on algorithmic choices. Two laboratories segmenting the same nuclei with different software, or the same software with different parameter settings, can obtain different counts, areas and classifications (7, 8).

Reviews have helped microscopists choose among the growing number of image-analysis packages (9–11), and community surveys and open-call applications record which packages researchers report using (12, 13). These surveys capture the practices of respondents who choose to participate. Community challenges have benchmarked algorithms on shared data for tasks such as cell tracking and single-molecule localisa-tion (14–17). Literature mining has counted software mentions in bioinformatics and across PubMed Central (18–20), and a scoping review mined MEDLINE for the image-analysis software used in cancer research (21). To our knowledge, no study has tracked mentions of image-analysis software across the full text of the microscopy literature over time. Over the past decade, deep-learning methods have entered routine microscopy image analysis. Cellpose introduced generalist cell segmentation, and StarDist introduced nucleus segmentation using star-convex polygons (7, 22). Throughout this study, we analysed how the software mentioned in microscopy papers changed as deep learning spread: whether mentions dispersed across many standalone packages or stayed with established platforms, how papers credit the packages they mention, and how they connect deep-learning methods to downstream measurements.

Here we present a census of 33 registry entries (Table S1) and a 2010-2025 count series for 32 entries across 1.7 million full-text microscopy papers (Fig. 1; Fig. 2). The registry covers 23 analysis packages, six microscope-control programs, MAT-LAB and Jupyter notebooks as references, Adobe Photoshop as image-editing software and NIH Image for historical comparison. The analysis aggregate contains the 23 packages; the 29-package total adds the six microscope-control programs.

**Fig. 1.**
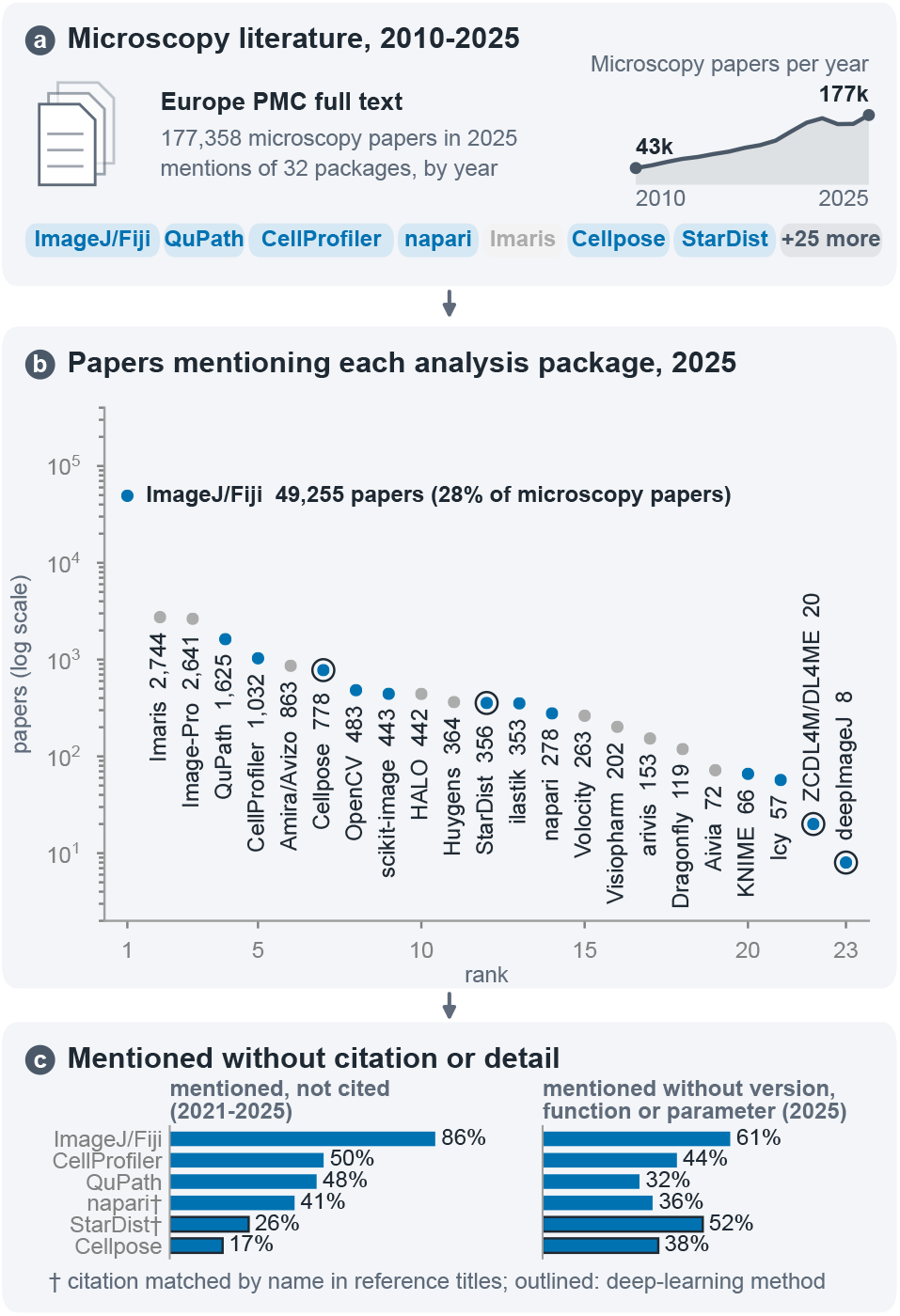
An overview of the publication landscape of image analysis packages. (**a**) We counted the full-text microscopy papers in Europe PMC that mention each of 32 registry entries, every year from 2010 to 2025. (**b**) Papers mentioning each of the 23 analysis packages in 2025, ranked on a logarithmic axis; blue marks open-source and grey commercial packages, and rings mark deep-learning methods. ImageJ/Fiji is mentioned in 28% of full-text microscopy papers. (**c**) For the six packages in the reporting audit, the share of papers mentioning or citing each package in 2021-2025 that mentioned it without citing its papers (Fig. 3; † marks napari and StarDist, whose own papers Europe PMC does not index, so their citations are matches of the package name in reference titles, Table S11), and the share of audited 2025 papers whose sentences mentioning it gave no version, function or parameter (Table S7); outlines mark deep-learning methods. ZCDL4M/DL4ME, ZeroCostDL4Mic/DL4MicEverywhere.

**Fig. 2.**
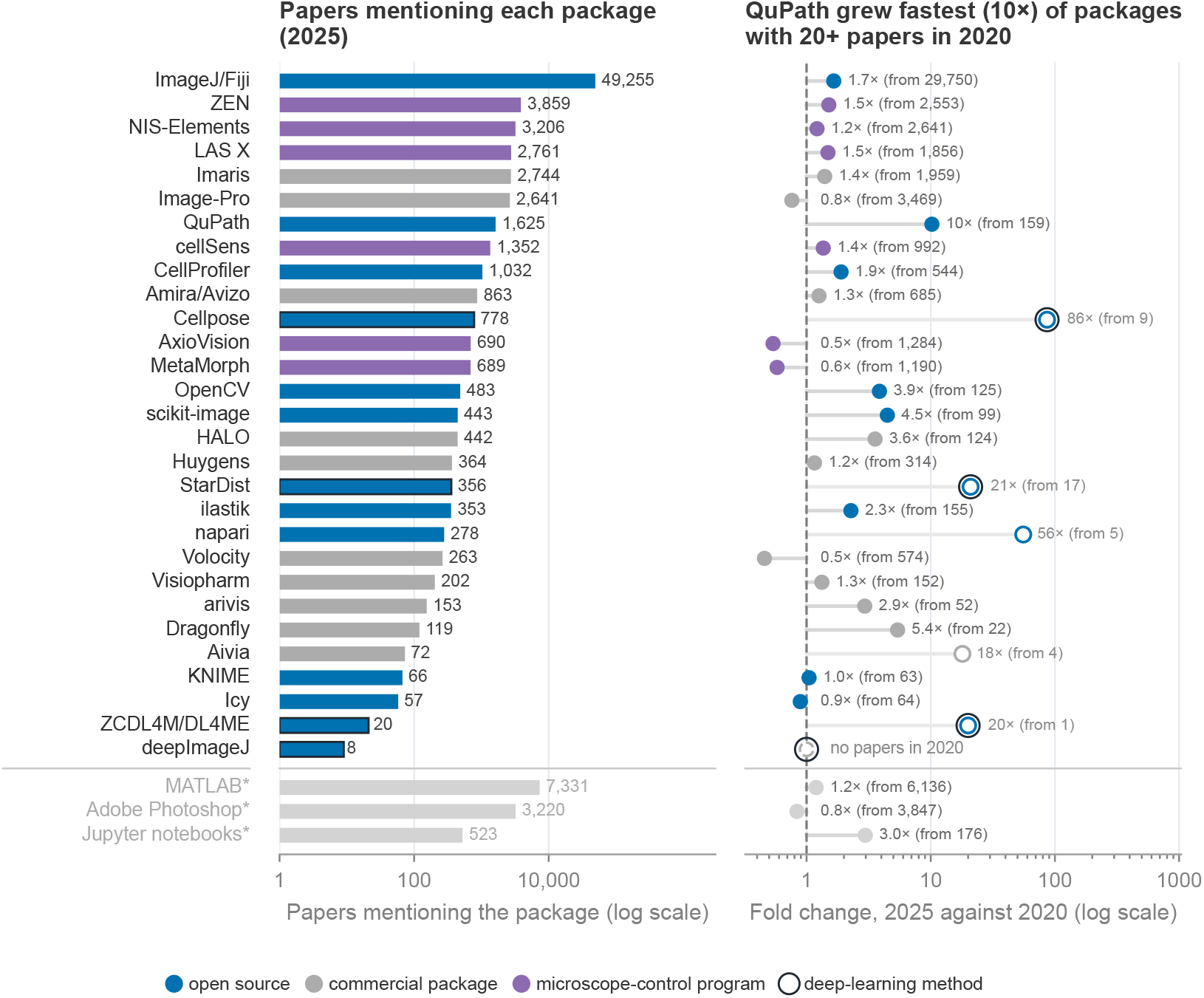
Papers mentioning each analysis package in 2025. Left, number of full-text Europe PMC microscopy papers mentioning each package in 2025 (logarithmic axis). Right, fold change in the annual number of papers mentioning each package between 2020 and 2025, labelled with the 2020 count, since a large fold change from a few papers is fragile; fold changes from fewer than 20 papers in 2020 are drawn as open markers with grey labels. The dashed line marks no change, and the open circle around it marks deepImageJ, which no indexed full-text microscopy paper mentioned in 2020 (preprint 2019; journal article 2021). Blue marks open-source packages, grey commercial packages and purple microscope-control programs (Methods; Table S1); a black outline marks the four deep-learning methods. MATLAB and Jupyter notebooks, marked with asterisks, are references for general numerical computing and for environments that run other packages; both appear separately for scale and enter no aggregate. Adobe Photoshop, also marked with an asterisk, is image-editing software and enters no aggregate. ZCDL4M/DL4ME, ZeroCostDL4Mic/DL4MicEverywhere.

MATLAB, Jupyter notebooks, Adobe Photoshop and NIH Image enter neither total. The NIH Image count series appears in Fig. S1; many mentions after about 2000 refer to ImageJ. For each entry in the 2010-2025 series, we measured how often papers mention it, how often they also cite its papers, which research areas those papers describe and which other packages they mention. We then read full-text samples of papers mentioning Cellpose, StarDist or ilastik together with ImageJ/Fiji to see how researchers combined them. Coding assistants may encourage narrow, purpose-built packages (23). These measurements provide a baseline for tracking whether such tools change the software that microscopy papers mention.

## Results

### The 33-entry registry and analysis totals

We tracked 33 registry entries (Table S1). Fifteen core registry entries span general image-analysis platforms (ImageJ/Fiji (24, 25), QuPath (26), CellProfiler (27), ilastik (28), napari (29), Icy (30) and KNIME (31)), deep-learning methods (Cellpose (7, 32), StarDist (22), deepImageJ (33) and ZeroCostDL4Mic with its successor DL4MicEverywhere, counted together as Zero-CostDL4Mic/DL4MicEverywhere (34, 35)), the commercial packages Imaris and arivis, and MATLAB and Jupyter note-books (36) as references for general numerical computing and for environments that run other packages. The other 18 entries include Image-Pro and NIH Image, seven other commercial analysis packages, the Python libraries scikit-image (37) and OpenCV (38), six microscope-control programs, and Adobe Photoshop as image-editing software. The six microscope-control programs enter only the 29-package aggregate that includes acquisition software. Adobe Photoshop, MATLAB, Jupyter notebooks and NIH Image enter neither aggregate. NIH Image stays outside the 2010-2025 comparison and appears in Fig. S1 and the supplementary tables; many mentions after about 2000 refer to ImageJ. For each entry in the 2010-2025 series and year, we counted the Europe PMC papers with indexed full text that mention the entry and microscopy (39), and divided by all such microscopy papers. To estimate research areas and deep-learning use, we read the title and abstract of every matching paper, or of a random sample when a package had more than 2,000 papers in a year (Methods). Fig. S1 extends the package-share series to 1995; the pre-2008 corpus is small and dominated by the back archives of a few journals.

Two language-model annotators checked the five existing vali-dation samples (Methods; Table S5). Names referred to software in 90-100% of checked matches for 30 of the 33 entries; Visiopharm (89%), Amira/Avizo (61%) and NIH Image (55%, 1995-2003 sample) fell below that range (Table S10). A single AI annotator labelled 40 full-text snippets for each new package. Image-Pro matched its software in 39 of 40 papers. All 40 Photoshop matches referred to Adobe Photoshop. Twenty-three papers used it on image data: 14 adjusted, cropped or converted images, 5 measured areas, counts or signal with it, and 4 did not say which. Eleven papers used it to assemble figures. NIH Image matched the historical program in 22 of 40 papers from 1995-2003 and in 15 of 40 papers from 2024-2025. After about 2000, “NIH Image” often refers to ImageJ. The deep-learning vocabulary had 94% precision and an estimated recall of 81%, and research-area keywords marked research in the area in 60% of checked matches. In 2025, 93% of ImageJ/Fiji matches mentioned the package in the Methods section (`methods_section_2025.csv` in the code repository). AI agents wrote the pipeline code and labelled the validation samples under our direction (Methods).

### Many papers mention a package without citing it

Citation counts are a common proxy for software use, but a paper can mention a package in its Methods and leave the package’s papers out of its references. We compared the two for the nine open-source packages whose own papers Europe PMC indexes, placing each paper as mentioning a package, citing its papers, or both (Methods). Of the 2021-2025 papers that mentioned ImageJ/Fiji or cited its papers, 86% mentioned it without citing them (Fig. 3a). The share was 48-50% for CellProfiler, QuPath and scikit-image and 9-31% for ilastik, Cellpose, deepImageJ and Zero-CostDL4Mic/DL4MicEverywhere. Preprints, which have no limit on references, mentioned ImageJ/Fiji without citing its papers less often than journal papers did (70% against 86%), while for the other packages the shares were similar (41% against 40%; Table S12).

**Fig. 3.**
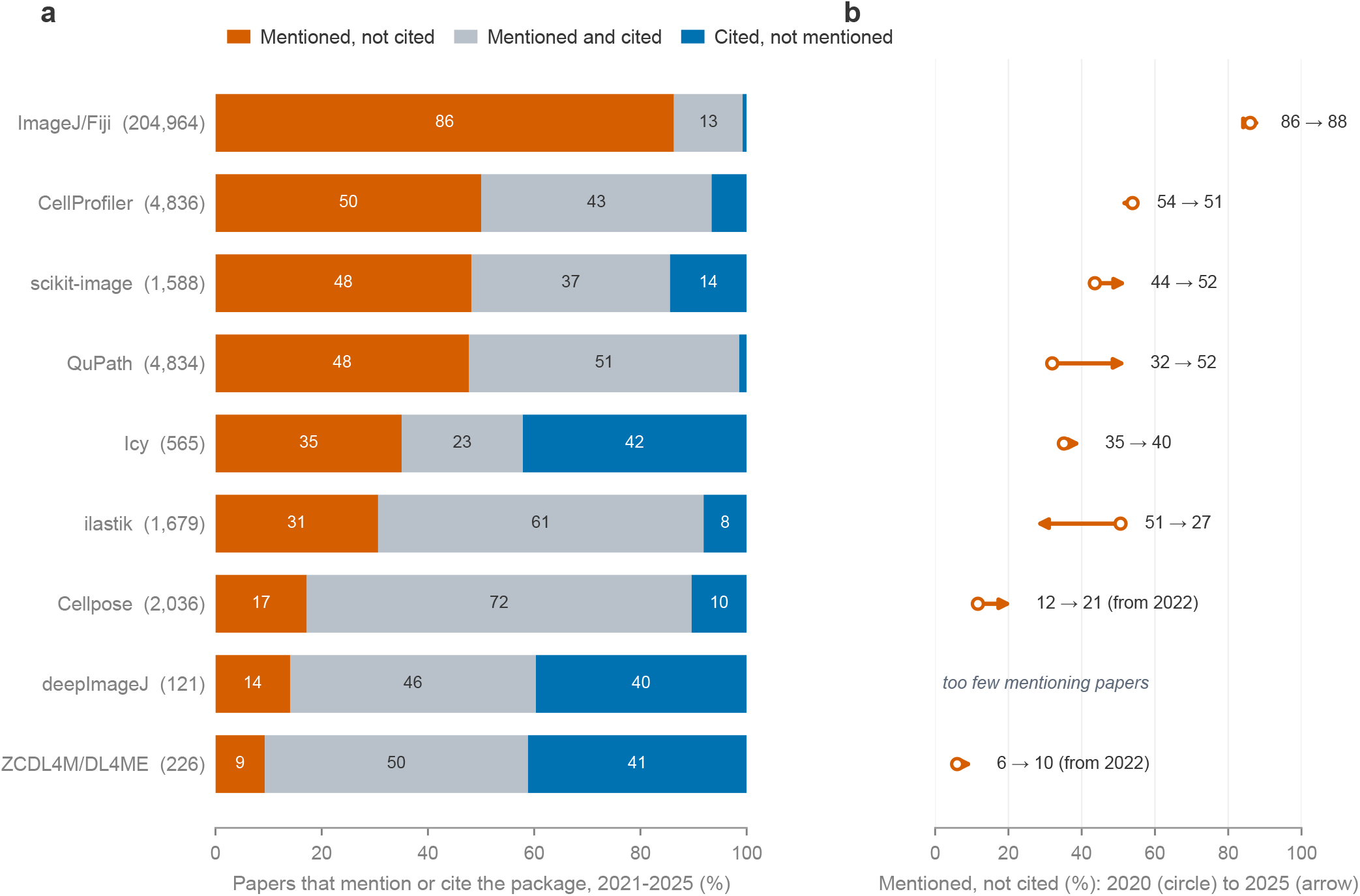
Many papers mention a package without citing it. For the nine open-source packages whose own papers Europe PMC indexes, full-text microscopy papers that mention the package or cite one of its papers through a citation link. (**a**) Papers in 2021-2025 that mention the package without citing its papers, do both, or cite it without mentioning it; the total is given beside each name, and rows are ordered by the first share. (**b**) Change in the mentioned, not cited share from 2020 (circle) to 2025 (arrowhead), in the rows of panel a; for Cellpose and ZCDL4M/DL4ME the start is 2022, the first year after their first paper with at least 20 mentioning papers. deepImageJ has fewer than 20 mentioning papers a year. Counts are as indexed, with no correction. Names carry a trailing wildcard, so mentioning counts exceed those of Fig. 2 (Table S9); Table S11 covers the five packages whose papers Europe PMC does not index. ZCDL4M/DL4ME, ZeroCostDL4Mic/DL4MicEverywhere.

For QuPath, the share rose from 32% in 2020 to 52% in 2025, and for Cellpose from 12% in 2022 to 21% (Fig. 3b). It stayed between 86% and 88% for ImageJ/Fiji and near half for CellProfiler, and fell from 51% to 27% for ilastik. Citation counts miss most papers that mention ImageJ/Fiji and a rising share of those mentioning QuPath and Cellpose.

The reverse case, a paper that cites a package without mentioning it, shows what our name queries miss. It was 14% or less for six packages and 40-42% for Icy, deepImageJ and Zero-CostDL4Mic/DL4MicEverywhere. Many papers citing these three contain none of the names our queries search for (for Icy, the query relies on product phrases), so mention counts understate them. Table S11 gives the five packages whose papers Europe PMC does not index, measured with reference titles.

### Mentions of analysis software grew faster than the microscopy literature

As the full-text microscopy literature grew about fourfold between 2010 and 2025, from 42,806 to 177,358 papers, the share of papers mentioning at least one of the 23 analysis packages rose from 15% to 31% (Table S2; Fig. 4). The share mentioning ImageJ/Fiji rose from 10% to 28%, and the share mentioning any of the other 22 packages rose from 5% to 7%.

**Fig. 4.**
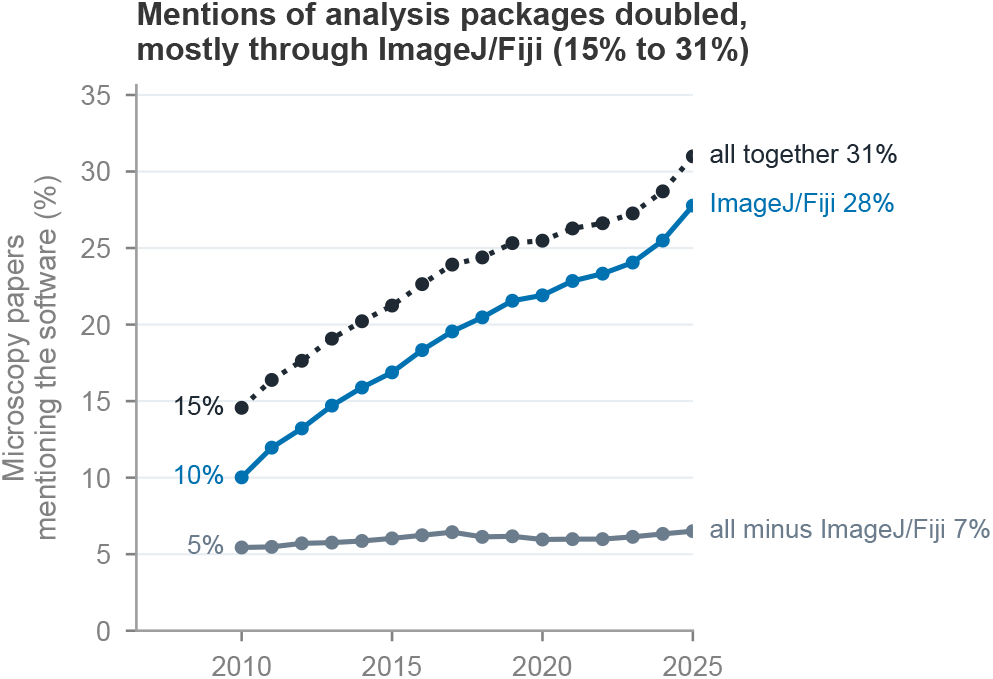
Mentions of analysis packages doubled, mostly through ImageJ/Fiji. Share of full-text Europe PMC microscopy papers mentioning at least one of the 23 analysis packages (all together, dotted), ImageJ/Fiji, and at least one of the 22 analysis packages other than ImageJ/Fiji (all minus ImageJ/Fiji). Each paper counts once per line, and Table S1 lists the packages. The number of full-text microscopy papers increased from 42,806 in 2010 to 177,358 in 2025.

Growth was uneven (Fig. 2; Fig. S1). CellProfiler rose 25-fold and ImageJ/Fiji about 12-fold. QuPath, first mentioned in 2017, reached 1,625 papers in 2025 and overtook CellProfiler in 2023. Image-Pro ranked third among analysis packages, with 2,641 papers in 2025. Older microscope-control programs and Volocity declined; ZEN, NIS-Elements and LAS X together accounted for 9,826 package-paper matches. From 2020 to 2025, the share of microscopy papers mentioning any of the other 22 packages rose from 6% to 7%; those packages accounted for 22% and 21% of analysis-package matches, respectively.

### Mentions concentrate on a few general-purpose platforms

As more packages came to be mentioned, a few platforms kept most of the mentions. The five packages most mentioned in 2025 accounted for 92% of analysis-package matches that year; the same five accounted for 89% in 2010. The largest, ImageJ/Fiji (24, 25), was mentioned in 28% of the full-text microscopy papers indexed in Europe PMC in 2025, up from 10% in 2010 (Fig. 4). Its share of package-paper matches in the 23 analysis packages rose from 67% in 2010 to 79% in 2025 (Table S2). These papers span cell counting, intensity measurement and treatment comparisons, and reach beyond cell biology into materials science (Fig. S2; Fig. 5). A mention count measures how widely a program is mentioned and favours general-purpose software used for routine steps; it does not rank programs by importance, quality or depth of analysis.

**Fig. 5.**
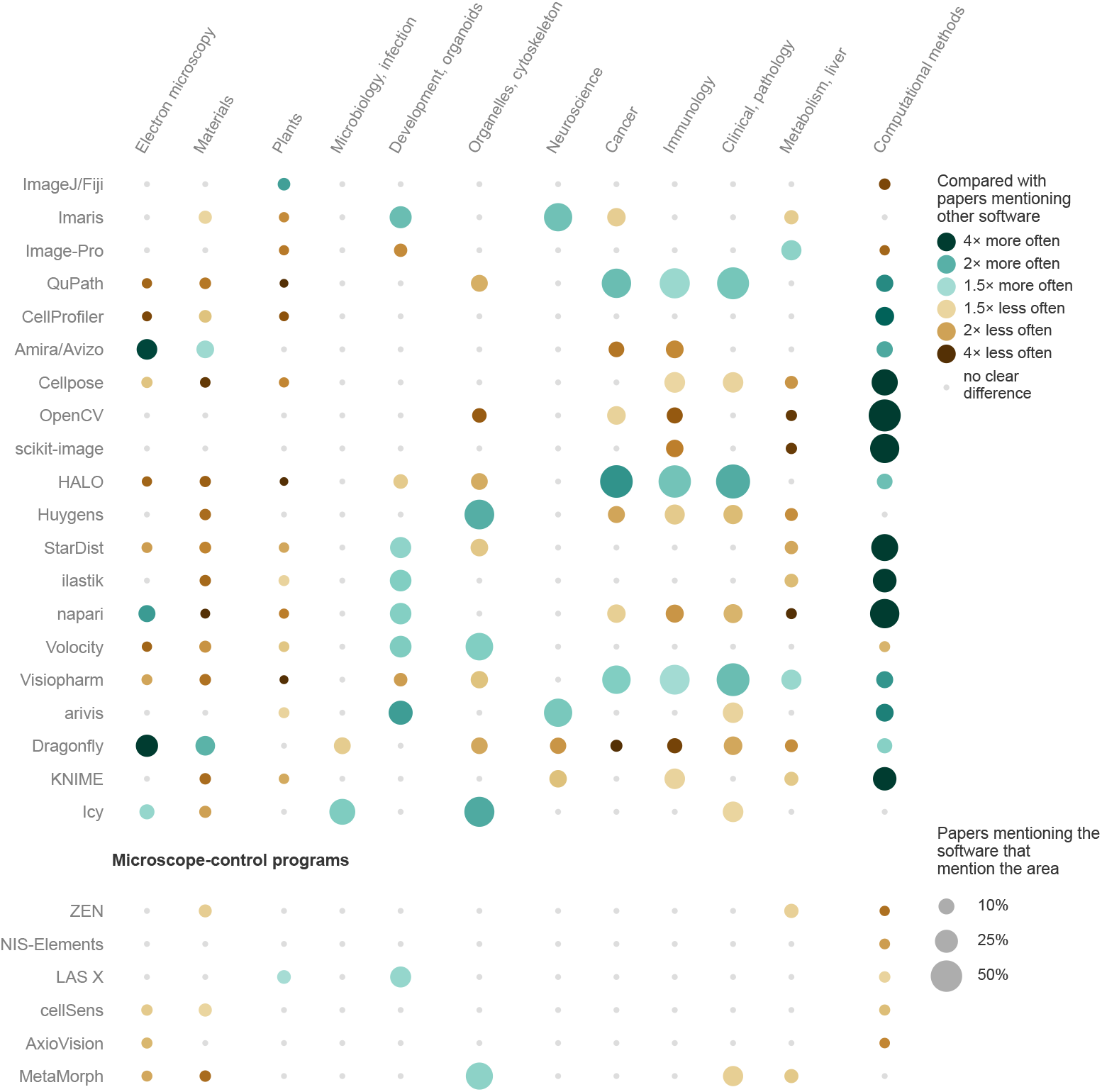
Research-area keywords associated with papers mentioning each program. For 2021-2025, a teal circle marks an area whose keywords appear in the titles or abstracts of papers mentioning the software at least 1.5 times as often as in those of papers mentioning other software, with a bootstrap 95% interval excluding equality (Methods); brown marks the corresponding depletion. Circle size indicates the weighted share of papers mentioning the software whose titles or abstracts contain the area’s keywords, and small grey dots mark pairs with no clear difference. We read every matching record except random samples of 1,000 per year for ImageJ/Fiji and Imaris and of 200 per year for Image-Pro. The comparisons are exploratory and do not adjust for multiple comparisons. A paper mentioning several programs contributes to each. Columns are grouped by topic (techniques and materials; organisms and development; cell biology; tissue and disease), and the final column gives computational-methods vocabulary. Areas whose keywords marked research in the area in fewer than half of checked matches are omitted (Methods). deepImageJ (62 records), ZeroCostDL4Mic/DL4MicEverywhere (114) and Aivia (178) fall below the 300-record display threshold. Each analysis package is compared with papers mentioning the other analysis packages. The lower block shows the six microscope-control programs, each compared with all papers mentioning the 23 analysis packages; for ZEN, NIS-Elements and LAS X we read random samples of 200 records per year. Fig. S2 gives associated terms and their prevalence.

Two sensitivity analyses test whether routine uses outside imaging drive these numbers. Counting only papers whose Methods mention both the package and microscopy, ImageJ/Fiji accounted for 80% of analysis-package mentions in 2025 (Methods). About half the microscopy papers mentioning ImageJ/Fiji also mention blots or densitometry, and removing every blot-mentioning paper leaves it with 62% of mentions in 2010 and 74% in 2025. With the six microscope-control programs included, its share rose from 43% to 66% between 2010 and 2025 (Table S2).

### Papers mentioning specialised platforms use the vocabulary of particular research areas

The areas measure vocabulary in titles and abstracts and agreed with MeSH headings, the subject terms that indexers assign to MEDLINE papers, for most displayed areas (Cohen’s kappa, an agreement score corrected for chance, 0.56-0.83), with the lowest agreement for clinical and pathology terms (agreement 12%, kappa 0.13; Table S6), which flag discussion of patients or diagnosis and do not establish a clinical study design.

QuPath was designed for digital pathology (26), where whole-slide images reach gigapixels and hold large numbers of cells, and the papers mentioning it carry the vocabulary of that field. They use clinical and pathology terms 1.8 times as often as papers mentioning other packages (Fig. 5), and they often concern tumours, immune microenvironments and patient cohorts. Within six years of its introduction, QuPath became the second most mentioned open-source analysis package after ImageJ/Fiji.

Papers mentioning Imaris and arivis use neuroscience terms 1.9 and 1.8 times as often, and arivis papers often describe light-sheet imaging of cleared tissue (Fig. S2). CellProfiler papers are associated with high-throughput screening (27), and ilastik papers, whose software trains classifiers interactively from user annotations (28), use developmental-biology and embryo terms 1.7 times as often. Papers mentioning napari, a multi-dimensional image viewer for Python (29), use electron-microscopy terms 2.2 times as often, as do papers mentioning Amira/Avizo and Dragonfly, at 3.7 and 4.3 times. Among commercial packages, HALO and Visiopharm papers use clinical and pathology terms 2.1 and 1.9 times as often.

The six microscope-control programs, each compared with all papers mentioning the 23 analysis packages, show fewer associations (Fig. 5). Papers mentioning LAS X use development and organoid terms 1.6 times and plant-biology terms 1.5 times as often, and papers mentioning MetaMorph use organelle and cytoskeleton terms 1.7 times as often. Computational-methods terms appear in the titles or abstracts of 2.0-3.3% of papers mentioning five of the six programs and of 4.4% of those mentioning MetaMorph, against 5.2% of papers mentioning analysis packages. Among analysis packages, the share is 3.4% for ImageJ/Fiji and 2.0% for Image-Pro, and it exceeds 30% for Cellpose, StarDist, napari, scikit-image and OpenCV.

These ratios compare each package with a baseline dominated by ImageJ/Fiji papers. They describe the vocabulary of the literature mentioning each package; software design, research practice, reporting and the composition of the indexed corpus may all contribute, and the analysis does not show why researchers chose a package.

### Papers mentioning the main deep-learning segmentation methods usually also mention an established platform

Two segmentation methods account for almost all mentions of the dedicated deep-learning methods we tracked: Cellpose (7, 32) and StarDist (22) took 98% of them in 2025. Their summed mentions rose from 1 to 20 per 1,000 mentions of the 13 core analysis packages between 2020 and 2025 (Fig. 6a). General architectures are mentioned more often: in 2025, 1,129 full-text microscopy papers mentioned U-Net, against 778 for Cellpose (Supp. Note 1).

**Fig. 6.**
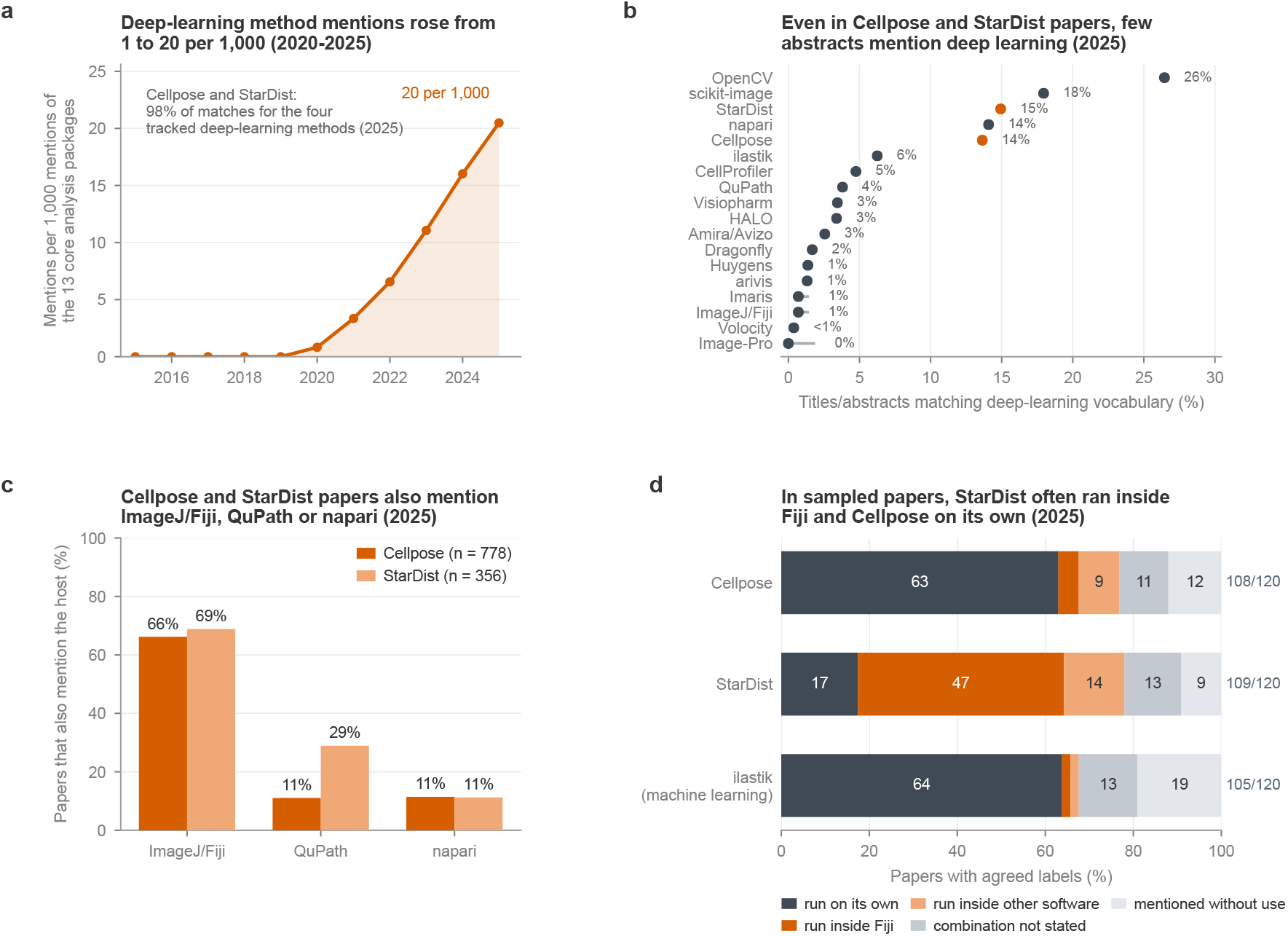
Deep-learning method mentions and how methods run alongside ImageJ/Fiji. (**a**) Summed package-paper matches for Cellpose, StarDist, Zero-CostDL4Mic/DL4MicEverywhere and deepImageJ, per 1,000 matches of the 13 core analysis packages (complete counts); Cellpose and StarDist account for 98% of matches for these four packages in 2025. (**b**) For analysis packages in the 23-package aggregate with at least 100 papers in 2025, the share of records whose title or abstract matches deep-learning vocabulary. Aivia (72 papers), KNIME (66), Icy (57), ZeroCostDL4Mic/DL4MicEverywhere (20) and deepImageJ (8) fall below that threshold. We read every matching record except random samples of 1,000 for ImageJ/Fiji and Imaris and of 200 for Image-Pro (Wilson 95% sampling intervals). Classification error remains in every estimate (Methods). (**c**) Share of 2025 full-text papers mentioning Cellpose (n = 778) or StarDist (n = 356) that also mention ImageJ/Fiji, QuPath or napari; a paper can mention more than one host (complete counts). (**d**) How Cellpose, StarDist and ilastik, an interactive machine-learning package whose Fiji plugin applies projects trained in the ilastik application, were run in three disjoint random samples of 40 open-access 2025 papers mentioning the method and ImageJ/Fiji: labels on which two language-model annotators agreed, pooled, as shares of the agreed papers (counts beside each bar). Dark bars indicate standalone execution, orange bars indicate execution inside a host, and grey bars indicate unspecified combinations or mentions without use. The labels cover papers mentioning both programs only.

In 2025, 66% of papers mentioning Cellpose and 69% of papers mentioning StarDist also mentioned ImageJ or Fiji; QuPath appeared in 29% of StarDist papers and 11% of Cellpose papers, and napari in 11% of each (Fig. 6c).

A co-mention does not show how researchers combined the packages, so two language-model annotators read the Methods of three random samples of 40 open-access papers mentioning both Cellpose and ImageJ/Fiji, and of samples of the same design for StarDist and for ilastik, an interactive machine-learning package that has relied mostly on random-forest classifiers (28) and now also runs deep-learning models from the BioImage Model Zoo (40) (Fig. 6d). The three methods were run differently. Papers most often ran Cellpose on its own: of the 108 papers on which the annotators agreed, 68 ran Cellpose on its own and described what they did with its output, 5 ran it inside Fiji, for example through TrackMate-Cellpose (41, 42), 10 inside other software, 12 did not say how the two were combined, and 13 mentioned Cellpose without using it. StarDist ran inside Fiji in 51 of 109 agreed papers and on its own in 19. Papers mentioning ilastik ran it on its own in 67 of 105 and inside Fiji in 2. ilastik’s Fiji plugin applies projects trained in the ilastik application, so every ilastik workflow involves the standalone program. Counting the 12 Cellpose papers on which the annotators disagreed as all standalone or as none bounds the standalone share (Methods). One author also read the full text of the first Cellpose sample, including linked repositories, and agreed with each model on 28 of 40 papers and with their consensus on 25 of 34. She found Cellpose run on its own in 19 of the 40 papers, inside other software in 8 and mentioned without use in 7, against 21, 4 and 4 of the 34 papers on which the models agreed, so the models may overstate standalone use. These labels describe papers mentioning both programs and cannot show how Cellpose users in general run it.

Hosts already integrate these methods. StarDist runs as a plugin in Fiji, QuPath and napari, and Cellpose through TrackMate-Cellpose, BIOP wrappers and deepImageJ (33). In 2025, 19% of papers mentioning StarDist and ImageJ/Fiji matched StarDist plugin phrases, against 4% of the corresponding Cellpose papers for Cellpose integration phrases; a plugin can also be used without being mentioned (Methods). Mentions of deepImageJ and Zero-CostDL4Mic/DL4MicEverywhere stayed below 30 papers a year.

Titles or abstracts matched deep-learning vocabulary in 14-15% of papers mentioning Cellpose or StarDist, and most often in papers mentioning the general Python libraries OpenCV (26%) and scikit-image (18%) (Fig. 6b). These shares measure explicit vocabulary, so differences in reporting and classifier error limit their reading as differences in deep-learning use.

### Papers mentioning Python packages more often describe computational methods

Computational-methods terms such as segmentation, pipeline and software appear four or more times as often in abstracts mentioning napari, Cellpose, StarDist, ilastik and KNIME as in abstracts mentioning other packages, and less often in abstracts mentioning ImageJ/Fiji (Fig. 5). Methods papers that build on these packages may explain the difference; the keyword analysis does not test it.

### The sentences mentioning analysis software often give no version, function or parameter

Readers cannot repeat an analysis from a package name, and imaging methods are often underreported (43). To see what papers report, we read the sentence mentioning each of six packages, and the sentence after it, in three random samples of 200 open-access 2025 papers for ImageJ/Fiji, QuPath and CellProfiler, and in every such paper for napari, Cellpose and StarDist (Methods; Table S7). Details given elsewhere in a paper are therefore not counted. These sentences gave no version, named function or parameter in 32% of QuPath papers, 36-44% of napari, Cellpose and CellProfiler papers, 52% of StarDist papers and 61% of ImageJ/Fiji papers, and restricting the audit to mentions in the Methods text changed no share by more than two percentage points. Papers mentioning napari or Cellpose shared code more often (78% and 58%) than papers mentioning ImageJ/Fiji (14%), although the shared code need not cover the audited analysis. Computational-methods vocabulary appeared in the titles and abstracts of few papers mentioning ImageJ/Fiji (4%, against 9-36% for other packages). Among papers without that vocabulary, those mentioning ImageJ/Fiji still gave only the name most often (61%), and stratifying by vocabulary changed the gap to the other packages from 21.9 to 21.5 percentage points (Table S8), so the difference in vocabulary explains little of the reporting gap. Within each package, papers with that vocabulary gave only the name about as often as the rest, within 6 percentage points, except for StarDist (41% against 56%).

## Discussion

We have shown that between 2010 and 2025, mentions of image-analysis software grew faster than the microscopy literature and stayed concentrated on a few general-purpose platforms. ImageJ/Fiji’s share of package-paper matches in the 23 analysis packages rose from 67% in 2010 to 79% in 2025, and 116 of 151 applications to the AI4Life open calls for image-analysis support mentioned ImageJ2/Fiji among the packages used or tested (13). From 2020 to 2025, papers mentioning any of the other 22 packages rose from 6% to 7% of the microscopy corpus; their share of analysis-package matches shifted from 22% to 21%. QuPath continued to gain mentions and appears more often in pathology papers. Napari papers more often use electron-microscopy terms, and Cellpose and StarDist are usually mentioned alongside an established platform. Older microscope-control programs lost mentions. Established platforms share features that may explain why researchers keep mentioning them, although this study does not test which features matter. Fiji installs as a self-contained application, runs on modest hardware without a graphics processor or an internet connection, and automates routine analyses through macros and plugins (44). QuPath opens gigapixel slides interactively and applies one script across a cohort (26), and CellProfiler pipelines, assembled without programming, run over large image sets (27). In napari, analysis code and the viewer for multi-dimensional images share one Python session (29). Python workflows require package installation and dedicated environments, and two methods in one analysis may need incompatible versions of the same library; LabConstrictor packages Python methods as desktop applications to reduce that cost for single methods (45).

Our counts measure mentions in records with indexed full text, and the Supplementary Information lists the study’s other limitations (Supp. Note 2). Mentions favour general-purpose programs used for routine steps and say little about which program carried out a paper’s central analysis, and a paper can mention a package without running it, as 13 of 108 papers mentioning both Cellpose and ImageJ/Fiji did. A mention also does not show where image processing ended and analysis began. Adobe Photoshop illustrates the problem. We kept it out of the analysis totals as image-editing software, yet 5 of 40 sampled papers measured vascular area, stained area or spot counts with its selection tools, and 4 more said only that they processed or analysed images in it. The same adjustments of brightness, contrast or colour count as analysis when a paper makes them in ImageJ/Fiji, so the boundary of the aggregate reflects our choice of registry as much as the authors’ practice. Microscope-control programs raise the same question from the acquisition side. ZEN and NIS-Elements include analysis modules, and Zeiss and Leica also sell arivis and Aivia, which our registry counts as analysis packages, so a paper that names acquisition software may also have analysed its images in it. In the microscope-control matches we checked, 90 of 162 papers used the program for analysis and 68 for acquisition only (Table S5). Vendor programs also run community methods. ZEN 3.14 denoises images with Noise2Void through the CAREamics library (46, 47), and arivis Pro segments cells with Cellpose (7, 48), so when a paper names only the vendor program our counts credit that program and miss the method. The vocabulary counts in Fig. 5 do not show which role a program played; that needs the workflow each paper reports. Changes in reporting, open-access coverage and corpus composition may affect the trends, and the corpus includes reviews and preprints as well as journal articles. The registry is a fixed selection, so its concentration estimates apply to the tracked packages (Supp. Note 1). Some packages cannot be counted by name: “CARE”, the content-aware image restoration method (49), matches 60,551 full-text microscopy papers in 2025, against 30 for its library name or full method name. Exact-name queries also miss papers in which the index joins a name to the citation number that follows it (“QuPath63”). A trailing wildcard would add 1% to the 2025 count of ImageJ/Fiji and 10-13% to those of Cellpose and StarDist, and would change ImageJ/Fiji’s share of analysis-package mentions by less than one percentage point (Table S9). Two language-model annotators labelled the validation samples, and a human annotator re-labelled only the first Cellpose sample.

Citation counts miss many of the papers that mention these packages. Across nine open-source packages, 9-86% of the papers that mentioned or cited a package in 2021-2025 mentioned it without citing its papers, and the share rose as QuPath and Cellpose spread (Fig. 1c; Fig. 3). Developers who estimate their number of users from citations have had to guess how many papers use a package for each paper that cites it, with plausible guesses from 1 to 5 (50). For papers that mention or cite a package, our counts put this ratio at 1.5 on average for eight packages and at 7.3 for ImageJ/Fiji. The ratio also changed within packages: between 2020 and 2025 it rose from 1.5 to 2.1 for QuPath and fell from 2.0 to 1.4 for ilastik, so a single guessed value misestimates both how many people use a package and how that number changes. We therefore propose counting the papers that mention or cite a package, *N*_*m*_, in full text and estimating user-paper occurrences as *N*_*u*_ ≈*F* × *P*× *U* × *N*_*m*_. Here *U* is the share of mentioning papers that ran the package, which our reading samples put at 0.87 on average for papers mentioning Cellpose, StarDist or ilastik together with ImageJ/Fiji; other packages and cite-only papers may differ. *F* counts the analysis projects for each one that reaches publication, and *P* the people who use the package in a paper; both still need usage logs or surveys. The product counts a person once for each paper, so a count of distinct users also needs the number of papers each user publishes with the package. Downloads can bound this estimate only within one distribution channel. QuPath’s installers were downloaded 131 times per citing paper in 2021-2025, within the range the image.sc discussion proposed, whereas Cellpose and StarDist, which notebooks and test pipelines reinstall, were installed from PyPI about 310 times per citing paper on average (Supp. Note 3).

The sentences that mention the software often lack details needed to repeat an analysis. In the reporting audit, they gave no version, function or parameter in 32-61% of papers, depending on the package, and only 14% of papers mentioning ImageJ/Fiji shared any code. Audits of imaging and statistical reporting have found similar gaps (43, 51). The QUAREP-LiMi initiative (52) has published community checklists for reporting image analyses at three levels (53). At the minimal level, a paper that uses an established workflow cites it, gives the versions of the workflow and of the software platform, reports key parameters, documents manual steps such as handdrawn regions of interest, and provides example input and output data. A paper that assembles a new workflow also gives the order of its steps and shares it as code or as a pipeline, and a paper that uses a machine-learning model makes the model and validation data available. Our audit checked three of these items, and only in the sentences next to each mention, so it cannot say how many papers meet the checklists elsewhere in their text. Standard image-quality metrics predict downstream analysis performance poorly (54), so a figure that looks good cannot show how its analysis was done. Papers that identify cell lines by research resource identifier report problematic lines less often (55), which suggests that a required reporting format changes what authors write. Journals and reviewers could ask for the QUAREP-LiMi minimal items in the same way, and Methods sections should also state how outputs passed from one program to the next.

Medina and Kornfeld argue that coding agents may encourage narrow, purpose-built packages and shift emphasis towards shared data formats (23). Dispersal would lower established platforms’ share of mentions and raise the number of newly mentioned packages; consolidation would preserve their share as new methods appear. Since 2020, ImageJ/Fiji’s share of mentions across the 23 analysis packages has held at 78% to 79% while its reach kept rising. These figures do not yet separate dispersal from consolidation. Separating them requires extracting software names beyond a fixed list, for example from software-mention datasets (18–20).

Foundation models are entering the same analysis workflows. Segment Anything for Microscopy runs as a napari plugin (56), Cellpose-SAM places the Segment Anything image encoder inside Cellpose (57), Cell-DINO describes cell morphology from fluorescence images without annotation (58), and SubCell, trained on the Human Protein Atlas images, describes protein localisation and cell morphology (59). Reproducing analyses with these models requires the model version, weights and inference settings (8), and the prompts for prompt-based models.

Whichever way the field moves, new methods have to work with the applications researchers already use. In our reading samples, papers most often ran Cellpose or ilastik on its own and passed the output to later steps. StarDist ran inside Fiji in about half of its papers. Six projects already link these applications. PyImageJ calls ImageJ from Python (60), napari-imagej runs ImageJ commands inside napari (61), deepImageJ runs trained models inside ImageJ (33), DL4MicEverywhere packages deep-learning notebooks in containers (35), the BioImage Model Zoo distributes models that deepImageJ, ilastik, QuPath and napari can read (40), and OME-Zarr provides a shared image and metadata format (6). A connection layer shared by these host applications could make a new method available in all of them at once. Its costs and benefits are untested, and standalone scripts and commercial workflows meet some of the same needs. The census provides a baseline for measuring whether such a layer is adopted.

## DATA AVAILABILITY

The processed counts, vocabulary estimates, sampled records, repository snapshots and annotator-labelled validation sets are available under CC BY 4.0 at https://github.com/HenriquesLab/mapping-bioimage-software (version 2026.10.31.2), with a Zenodo archive at https://doi.org/10.5281/zenodo.23844634. The repository includes publication identifiers and no article text; release/refetch_identifiers.py rebuilds the text corpus from the identifiers through Europe PMC. Later queries to public endpoints may return updated records.

## CODE AVAILABILITY

The analysis and figure code is available under the MIT licence at https://github.com/HenriquesLab/mapping-bioimage-software (version 2026.10.31.2), with a Zenodo archive at https://doi.org/10.5281/zenodo.23844634. File names given in Methods refer to its census package or release folder.

## AUTHOR CONTRIBUTIONS

**Conceptualisation:** T.S., B.M.S., G.J., R.H. **Methodology:** T.S., B.M.S., I.C., G.J., R.H. **Software:** T.S., B.M.S., R.H. **Validation:** T.S., B.M.S., I.C., R.H. **Formal analysis:** T.S., B.M.S., G.J., R.H. **Data curation:** T.S., R.H. **Visualisation:** T.S., B.M.S., R.H. **Writing - original draft:** T.S., B.M.S., R.H. **Writing - review and editing:** T.S., B.M.S., I.C., C.J., G.J., R.H. **Supervision:** C.J., G.J., R.H. **Funding acquisition:** C.J., G.J., R.H. T.S. and B.M.S. contributed equally and are co-first authors.

## ACKNOWLEDGEMENTS

We thank the developers and communities behind the open-source bioimage-analysis packages analysed here. We thank George McNamara for suggesting the pre-2010 extension and the additional packages, and Sebastian Rhode (Carl Zeiss Microscopy) for asking why ZEN was absent from the research-area comparison, which led to the microscope-control block in Fig. 5.

## EXTENDED AUTHOR INFORMATION

## Methods

### Package registry and queries

The registry (`tools.yml`) defines 33 entries (Table S1): 23 analysis packages, six microscope-control programs, two references (MATLAB and Jupyter notebooks), Adobe Photoshop as image-editing software and NIH Image for historical comparison. A package’s query joins its aliases with OR and adds `AND (microscopy OR microscope) AND HAS_FT:Y`, and the yearly query adds `AND PUB_YEAR:<year>`. Icy, Huygens, HALO, ZEN and Dragonfly use product phrases or vendor names because their bare names are ordinary words. The NIH Image query excludes records that mention ImageJ under any of six spellings, so it also drops NIH Image papers that mention ImageJ. The authors chose the registry to cover each kind of image-analysis software, so it is a purposive selection (Supp. Note 1 gives the rule for adding packages). The 15 core entries comprise 13 analysis packages and the two references, MATLAB and Jupyter notebooks; the 13 analysis packages form the denominator of the deep-learning series (Fig. 6a). Eighteen further entries include Image-Pro, NIH Image, seven other commercial analysis packages, two Python libraries, six microscope-control programs and Adobe Photoshop as image-editing software. The 23 analysis packages form the analysis aggregate; the 29-package total adds the six microscope-control programs. Adobe Photoshop, MATLAB, Jupyter notebooks and NIH Image enter neither aggregate. The NIH Image count series appears in Fig. S1; many mentions after about 2000 refer to ImageJ. MATLAB and Jupyter notebooks run other packages within them, so counting them beside dedicated analysis packages would count some analyses twice. In the microscope-control matches we checked, 68 of 162 described acquisition only, so we report concentration with and without these programs. Two authors (G.J. and R.H.) developed ZeroCostDL4Mic and DL4MicEverywhere.

ImageJ (24), its rewritten core ImageJ2 (44) and the Fiji distribution (25) form one ecosystem, so the registry counts “ImageJ” OR “ImageJ2” OR “Fiji” as one package. In 2025, 90% of these papers mentioned ImageJ and 25% mentioned Fiji. Because Fiji is also a country, we read 40 random titles of 2025 Fiji-only matches; all were bioimaging studies, and fewer than 1% of Fiji-only papers contained country terms such as Suva or Fijian.

### Literature and repository data

Europe PMC (39) indexes full text alongside titles and abstracts. We restricted every query and the microscopy denominator to records with indexed full text (`HAS_FT:Y`); abstractonly records fell from 34% of microscopy records in 2010 to 11% in 2025. The queries impose no publication-type restriction, so reviews and preprints contribute, and a preprint and its journal version count separately. Yearly OpenAlex counts (63), which cover titles and abstracts only, provide a cross-check (Table S2). We collected the original 2010-2025 Europe PMC and OpenAlex counts and the deep-learning sample on 2 October 2026. On 28 October 2026, we queried the three added packages in Europe PMC and OpenAlex, extended the Europe PMC series back to 1995, refreshed the host co-mentions and query-based sensitivity analyses, and drew the added packages’ abstract samples. Europe PMC counts drift slightly as records are added. The forum.image.sc site blocks automated collection and contributes no data.

Reach divides a package’s yearly matches by all indexed full-text microscopy records that year. Concentration divides them by the summed matches of the 23 analysis packages, or of the 29-package aggregate when the six microscope-control programs are included. Adobe Photoshop, MATLAB, Jupyter notebooks and NIH Image enter neither aggregate. A record mentioning two or more packages contributes once to each, so sums count package-paper matches.

### Deep-learning estimates

For each package and year, we retrieved every matching identifier and read the title and abstract of every record. Where a package-year exceeded 2,000 matches, we read a seeded simple random sample: 1,000 records for ImageJ/Fiji in every year and for Imaris in 2021-2025, and 200 for Image-Pro, MAT-LAB and the largest microscope-control package-years. We avoided Europe PMC’s relevance order, which over-samples papers that feature a package prominently: for Cellpose in 2025, the first 200 records give a deep-learning share of 35%, more than twice the share among all papers. A record counts as deep learning if its title or abstract matches a lexicon term and no exclusion guard applies; terms match on word boundaries and tolerate hyphen and space variants. Sampled package-years carry Wilson 95% intervals (Table S3), and package-years read in full carry none. The field-level share is the count-weighted mean of the core packages’ shares, with a finite-population-corrected interval.

As a check, a second estimate queries Europe PMC for ImageJ/Fiji, napari and Jupyter notebooks together with a deep-learning term from `dl_in_tools_lexicon.yml` in the title or abstract, excluding reviews and preprints from numerator and denominator alike. In 2025 it gives 0.5% for ImageJ/Fiji, 12% for napari and 12% for Jupyter notebooks, close to the lexicon estimates in Table S3. The mentioned deep-learning series sums the full-text matches of Cellpose, StarDist, deepImageJ and Zero-CostDL4Mic/DL4MicEverywhere.

*Lexicon, version 2*. Include terms: deep learning; deep neural network; convolutional neural network; convolutional network; neural network; CNN (case-sensitive); U-Net; UNet; transformer; vision transformer; self-supervised; semantic segmentation; instance segmentation; generative adversarial; GAN (case-sensitive); diffusion model; foundation model; encoder-decoder; ResNet; VGG (case-sensitive); autoencoder; representation learning; pretrained model; pre-trained model; segmentation model; denoising network; Noise2Noise; Noise2Void; CARE (case-sensitive); content-aware restoration; attention mechanism. Exclusion guards: without deep learning; non-deep-learning; classical machine learning only; no neural network; compiled standalone application. Version 2 matches CNN, GAN, VGG and CARE case-sensitively, which stops gene symbols and capitalised words from matching.

### Host co-mentions and the Cellpose sample

For Cellpose and StarDist in each year from 2020 to 2025, we counted full-text microscopy records mentioning the method and records also mentioning ImageJ/Fiji, QuPath or napari (`build_hosts.py`). Within ImageJ/Fiji co-mentions, we counted integration phrases such as StarDist plugin, TrackMate-StarDist, Cellpose plugin or wrapper, TrackMate-Cellpose, BIOP and deepImageJ; these phrases count named integrations only. From the open-access 2025 papers mentioning both Cellpose and ImageJ/Fiji, we drew three disjoint seeded samples of 40, and two annotators read each paper’s Methods and labelled how Cellpose was run (Validation samples). We repeated the design for StarDist and ilastik, from frames of 200 and 215 papers, with the same question and the method’s name and Fiji integrations substituted (`workflow_hosts.py`).

### Research areas

We reused the records read for the deep-learning estimate for 2021-2025 and the 23 analysis packages: 39,502 package-paper records covering 34,222 distinct papers, each weighted by the inverse of its package-year sampling fraction. For the six microscope-control programs we used their 18,541 package-paper records, read in the same way, and compared each program with the analysis-package records, which leaves the analysis baseline unchanged (`control_enrichment` in `domains.py`). Patterns in `domains.py` match thirteen research areas and a computational-methods vocabulary in titles and abstracts. We compared keyword areas with MeSH descriptors mapped per area in a stratified sample of 3,000 MED-source papers, 2,237 with MeSH headings, using `mesh_check.py` (Table S6). Two independent annotators checked three samples of 8 keyword matches per area. For the 11 areas in Fig. 5, 52-100% of agreed matches marked research in the area; we omit cardiovascular and renal (39%) and single-cell and spatial omics (45%), which fell below half. Electron microscopy is scored by use of the technique (24 of 24 agreed matches). Enrichment divides a package’s weighted keyword share by that of the records mentioning the other analysis packages; this baseline is dominated by ImageJ/Fiji in every row except ImageJ/Fiji’s own. Intervals come from 500 Poisson bootstrap replicates that give each sampled paper one weight across every package it appears under and hold fully read strata fixed. They describe sampling stability, exclude classification error and are not adjusted for multiple comparisons. Associated words (Fig. S2) are ranked by log-odds against the same baseline, counted once per abstract and restricted to words present in at least 2% of a package’s abstracts.

### Reporting audit

For each of six packages, we queried Europe PMC for the package’s aliases in the METHODS field with the registry context, `OPEN_ACCESS:Y` and `PUB_YEAR:2025`. The frames held 41,322 ImageJ/Fiji, 1,251 QuPath, 764 CellProfiler, 179 napari, 517 Cellpose and 244 StarDist papers. We drew three disjoint seeded samples of 200 papers from the first three frames and read every paper of the last three, divided at random into three replicates (`replicates.py`). For each paper we took the sentence mentioning the package and the sentence after it, from Methods sections located by section type or heading, or from the body text where the name appeared only there (139 of 2,737 papers with full text). We scored four indicators with fixed text patterns: a version number next to the name; a named plugin, command, module or model; a parameter value, named thresholding method or statement of default settings; and, anywhere in the article, shared code, meaning a code-hosting link, an analysis file or a statement that code is available, but not code available on request. Sampled packages carry Wilson 95% intervals, and packages read in full carry none. Two annotators labelled three samples of 60 papers (10 per package) from the same text without seeing the pattern results. They agreed on 92-100% of labels per indicator (Cohen’s kappa 0.80-1.00), and against their shared labels the patterns reached 92% precision for version, 81% for function, 78% for parameter and 84% for code, with recall of 97%, 72%, 75% and 92%. We did not adjust the patterns after seeing these labels. Across the three ImageJ/Fiji samples, the name-only share ranged from 56% to 67%. We used `audit_by_area.py` to flag the Figure 5 computational-methods pattern (segmentation, deep learning, machine learning, neural networks, pipelines, open-source, software, algorithms or computational) in audited papers’ titles and abstracts and compared name-only shares with a vocabulary-stratified Mantel-Haenszel risk difference (Table S8).

### Mentions and citations

For each open-source package and year, we counted full-text microscopy papers that mention the package, papers that cite it, and papers that do both (`build_citations.py`). A paper cites a package when Europe PMC links one of its references, through the `CITES` field, to one of the package’s own papers, the papers its developers ask users to cite (`package_papers.yml`). This works for the nine packages whose own papers Europe PMC indexes. The papers of napari (a Zenodo software record), StarDist (conference proceedings), Jupyter, KNIME and OpenCV are not indexed, so for these packages we report matches of the package name in reference titles, which also admit plugin papers mentioned after a host (Table S11). Names carry a trailing wildcard, because the index joins a name to a citation number that follows it (Table S9). Trends start in the year after a package’s first own paper, when citing it first became possible. Counts are as indexed and carry no correction; samples of papers read in full text measured how often the index missed a name or citation (Table S10). To compare preprints (Europe PMC source PPR) with journal papers, we restricted both to records whose reference list Europe PMC has parsed (`HAS_REFLIST:Y`), because it has parsed 60% of full-text microscopy preprints and 99% of journal papers (`build_citations_by_source.py`).

### Validation samples

We checked the text queries with five samples, each labelled by two annotators (Table S5). For each, `replicates.py` drew three disjoint seeded samples from a recorded frame, and two language-model annotators labelled every item without seeing the query or classifier result (`label_replicates.py`, `label_round2.py`). Estimates use the labels on which both annotators agreed. They disagreed on 10 of 450 alias items for the added packages, 1 of 413 for the core packages, 30 of 600 abstracts, 39 of 336 keyword matches, 12 of 120 Cellpose papers, 11 of 120 StarDist papers and 15 of 120 ilastik papers. The samples contained ten random 2025 matches per package, labelled for whether the name referred to the software and, for microscope-control programs, whether it concerned acquisition or analysis; 200 abstracts per replicate stratified by lexicon result; 8 keyword matches per research area; and 40 open-access 2025 papers mentioning ImageJ/Fiji together with Cellpose, StarDist or ilastik. Counting the 12 disagreements as all standalone or as none, the standalone share of the 120 papers lies between 57% and 67%. Of these disagreements, 7 were items the GLM-5.3-Flash annotator left unlabelled: 1 alias item for the added packages, 2 abstracts and 4 keyword matches.

For the three new registry entries, a single AI annotator labelled 40 Methods or full-text snippets per entry. The Image-Pro and Photoshop samples came from 2025; the NIH Image sample came from 2024-2025. We classified the recent NIH Image matches by meaning and did not use them to estimate precision for the historical program. A separate sample of 40 full-text matches from 1995-2003 estimated historical-program precision; only clear references to that program counted as software. An independent review checked the 2025 labels. Two language-model annotators then separately relabelled the 23 Photoshop papers that used it on image data as adjustment, measurement or unclear; they agreed on 22, and the authors resolved the remaining paper.

The deep-learning lexicon was correct for 215 of 228 agreed lexicon-positive abstracts and missed 4 of 342 agreed lexicon-negative abstracts. The frozen validation sample includes Jupyter notebooks; the aggregate population excludes both reference packages. Scaling these error rates to the 24,979 records in the core-package aggregate (1,236 lexicon-positive) gives a precision of 94% and an estimated recall of 81% (95% interval 62-92%, with precision held fixed). These pooled estimates do not give separate error rates for each package.

In the second labelling round, research-area keywords marked research in the area in 178 of 297 agreed matches (60%), from 39% for cardiovascular and renal to 89% for computational methods; Fig. 5 omits the two areas below 50%. Electron microscopy is defined by use of the technique, but the second-round question counted a keyword used as a method as a passing match. We therefore relabelled its 24 items with a question asking whether the study uses or develops electron microscopy, and the annotators agreed that all 24 did.

### Sensitivity analyses

We repeated the yearly queries requiring the package and microscopy in Methods, with the same restriction on the denominator (`build_sensitivity.py`; Table S4). ImageJ/Fiji reaches 12% of these papers in 2010 and 33% in 2025, and takes 67%, 78% and 80% of analysis-package men-tions in 2010, 2020 and 2025. This query retains 90% of ImageJ/Fiji matches and 76-79% of napari, Cellpose and StarDist matches in 2025 (Table S4). It cannot separate microscopy analysis from other assays described in the same Methods. For blots, we split each package’s matches by whether the article also mentions western blot, immunoblot, blotting, densitometry, densitometric or gel quantification (`build_blot_sensitivity.py`). A paper can mention blots and still analyse microscopy in ImageJ/Fiji, so removing blot-mentioning papers bounds their contribution from above. In 2025, 53% of microscopy papers mentioning ImageJ/Fiji also mentioned blots or densitometry. A trailing wildcard on single-word aliases measures the exact-name undercount (Table S9).

### Use of AI assistants

AI agents wrote the analysis code, labelled the validation samples and curated the literature for this study under the authors’ direction between July and October 2026.

*Language models*. Anthropic’s Claude (Opus 4.8, Opus 5 and Opus 5.5, with Sonnet 5.5 for some delegated tasks) ran most of these tasks. OpenAI’s GPT-5.6 and GPT-6 (Terra, Luna and Sol) and Google’s Gemini Flash (versions 3.6 to 3.8) ran delegated coding, review and annotation tasks. Open-weight models labelled most validation samples: DeepSeek-V4.1-Flash and GLM-5.3-Flash, served by Synthetic (https://synthetic.new), were the two independent annotators, and Synthetic’s routed `syn:large` alias, whose underlying model was not recorded, labelled the reporting audit with Gemini 3.8 Flash (`data/validation/ANNOTATORS.md`).

GLM-5.2, served through OpenRouter, ran some delegated tasks, and an open-weight model on a local GPU server reviewed draft prose for style. The models in use changed during the project as providers released new versions.

*Agent programs*. The models worked through two agent programs, which let a language model read and write files, run code and call other tools. Pi (https://pi.dev; https://github.com/earendil-works/pi), an open-source terminal agent that sends each task to a chosen model, did most of the work, and Claude Code (https://github.com/anthropics/claude-code), Anthropic’s agent, revised figure code and retrieved literature.

*Tasks*. The agents wrote and debugged the Python modules that query Europe PMC, OpenAlex, GitHub and PyPI, draw the samples, compute the estimates and draw the figures. They assigned the labels in the validation samples, recording the phrase each label rests on. They curated the literature: they checked each reference’s DOI, title and authors against Crossref and Europe PMC, downloaded open-access PDFs of cited works, and recorded each in a lab reference library with its DOI and abstract, so they could check the claim attached to a citation against the paper. Most PDFs came from direct downloads; for bioRxiv and PubMed Central, the agents downloaded PDFs through a browser with OpenCLI (https://github.com/jackwener/OpenCLI), an open-source command-line program that drives a signed-in browser, and flagged paywalled papers for an author to fetch by hand. The agents also drafted and revised passages of the manuscript, which the authors edited and approved.

*Oversight and records*. The authors chose the packages, wrote the search queries and the deep-learning and research-area vocabularies, specified the analyses, and checked the outputs against the data before accepting a change. Each agent program kept session logs recording which model did which task; these logs do not cover assistants used on co-authors’ own computers. Two language-model annotators labelled the five paired-annotator samples independently. A single AI annotator labelled 40 papers for each of the three new registry entries, and an independent review checked the 2025 labels. The historical NIH Image sample was coded separately. One author (T.S.) labelled the 40 papers of the first Cellpose sample by hand from the full text, on a blank sheet that did not show the model labels, with the first-round rubric; its three standalone labels merge into one for comparison (`human_check.py`).

### Statistics and reproducibility

Counts enumerate matches to the stated Europe PMC queries, so package-years read in full carry no sampling error; classification error remains in every estimate. Wilson intervals describe sampling error only. Every sample records its seed and frame (`replicates.py`). We ran no hypothesis tests. Live Europe PMC additions and record changes can alter rerun results, so exact reproduction requires the archived data. The pipeline used Python 3.11.16 with requests 2.32.4, pandas 2.3.0, numpy 2.3.1, matplotlib 3.11.2 and PyYAML 6.0.2.

## Supplementary Information

### Supp. Note 1: Packages and methods outside the registry

The registry omits Fiji plugins mentioned on their own, the data-management platform OMERO and general programming languages, and it tracks only four dedicated deep-learning methods. The 2025 full-text microscopy query finds 1,129 papers mentioning U-Net and 155 mentioning Segment Anything (`build_sensitivity.py`). U-Net is an architecture inside many packages, Cellpose included, so such counts overlap with the registry. Cellpose-SAM (57) is mentioned in 17 papers, all of which also mention Cellpose, and Segment Anything for Microscopy (56) in 15. Among packages without deep learning, TrackMate (41, 42) is mentioned in 506 papers and Trainable Weka Segmentation (64) in 132; 95% and 98% of these also mention ImageJ or Fiji and so fall within its count.

We will extend the registry by a fixed rule: a package enters when an independent source mentions it, it has a paper and its name can be counted, however often it is mentioned. Software that applicants to the AI4Life open calls mentioned, and that the resulting pipelines used (13), supplies the first candidates: in 2025, Labkit (65) is mentioned in 79 papers, BigWarp (66) in 39 and Noise2Void (46) in 29.

### Supp. Note 2: Limitations of the study

#### What is measured

Every count is a mention: a paper whose indexed full text contains the program’s name. A mention does not show that the program ran, which step it performed or how central that step was. Mentions favour general-purpose programs used for routine steps such as counting, intensity measurement or blot densitometry, so the counts describe how widely software is mentioned and do not rank programs by importance, quality or depth of analysis. Restricting matches to Methods preserves the trends but cannot separate microscopy analysis from other assays in the same Methods.

#### Corpus

The counts come from Europe PMC records with indexed full text, which over-represent open-access journals and change in coverage over time. The microscopy filter is a word match, which can admit papers whose microscopy is incidental and miss papers that use other terms. The pre-2008 corpus in Fig. S1 is small and dominated by a few journals’ back archives; the shaded years provide historical context. Europe PMC counts drift as records are added, so the archived snapshot is the reference.

#### Registry and queries

The 33 registry entries are a purposive selection. Concentration uses the 23 analysis packages or the 29-package total that adds microscope-control programs; software outside the registry would lower both shares (Supp. Note 1). Adobe Photoshop is recorded as image-editing software, and NIH Image stays outside the 2010-2025 comparison and appears in Fig. S1 and the supplementary tables; neither enters an aggregate. Queries miss papers that describe a program without mentioning it. Icy, ZEN, HALO and Dragonfly are ordinary words, Amira a personal name and Fiji a country, so their queries use product phrases, vendor names or a country check, trading missed mentions for precision (Table S10). Some packages cannot be counted by name: “CARE”, the content-aware image restoration method (49), matches 60,551 full-text microscopy papers in 2025, while its library name CSBDeep or its full name matches 30, a lower bound on its use; the abbreviation N2V for Noise2Void is similarly ambiguous. Exact names also miss papers in which the index joins a name to the citation number that follows it (“QuPath63”). The undercount is larger for newer packages: a trailing wildcard adds 1% to ImageJ/Fiji’s 2025 count, 3-5% to QuPath and CellProfiler and 7-13% to the Python libraries, napari, ilastik, Cellpose and StarDist, so ImageJ/Fiji’s share of analysis-package mentions is slightly overstated, 78.7% against 78.3% with the wildcard, with no change of rank (Table S9). Papers that cite a package without mentioning it give a second estimate of these misses (Fig. 3a).

#### Citations

A citation counts when Europe PMC links a reference to one of a package’s own papers. References that Europe PMC did not parse or link go uncounted, and five packages whose papers it does not index are measured only by reference titles (Table S11).

#### Classification and labels

The deep-learning vocabulary reads titles and abstracts, so it misses uses that papers do not name. For the displayed research areas, keywords matched 52–100% of checked papers (Methods). These enrichment results describe vocabulary patterns in the literature. We did not test why researchers chose a package.

Two language-model annotators independently labelled the five paired-annotator samples. Estimates use items on which both agreed. Disagreement rates ranged from under 1% to 13% by sample.

A single AI annotator labelled 40 full-text/Methods snippets for each new package, plus a separate 40-paper historical sample for NIH Image. The resulting false-positive rates appear in Table S10. The workflow samples cover papers mentioning ImageJ/Fiji with Cellpose, StarDist or ilastik.

#### Reporting audit

The audit reads the sentence mentioning a package and the sentence after it, so details given elsewhere in a paper are missed. Code sharing counts anywhere in the article and need not cover the audited analysis.

#### Scope

Download counts include automated testing and installation as a dependency. The study does not measure package accuracy, usability, cost or the reasons behind software choice.

### Supp. Note 3: Estimating users from citations and downloads

An image.sc discussion proposed the formula *N*_*u*_ = *F* × *C*× *P* × *N*_*c*_ to estimate how many people use a package (50). *N*_*c*_ counts the papers that cite the package, and *C* the papers that use it for each paper that cites it. *F* counts the analysis projects for each one that reaches publication, and *P* the people who use the package in a paper. Contributors guessed *F* at 3-10, *C* most plausibly at 1-5 and *P* at about 3, which puts the product *F* × *C* × *P* between 9 and 150.

#### Measured

*C*. Our full-text counts measure *C* as the papers that mention or cite a package for each paper that cites it: 1.5 on average for eight packages and 7.3 for ImageJ/Fiji in 2021-2025 (Table S11). The only survey-based value in the discussion, about 1.3 for QuPath from a review of the papers using it, lies close to our 1.5 for 2020. *C* also changed within packages: between 2020 and 2025 it rose from 1.5 to 2.1 for QuPath and fell from 2.0 to 1.4 for ilastik, while it rose from 7.1 to 8.1 for ImageJ/Fiji and stayed near 2 for CellProfiler. A single guessed *C* misestimates both the number of users and its change over time.

#### Mentions and use

A mention does not show that a paper ran the package. Our workflow samples read 2025 papers mentioning Cellpose, StarDist and ilastik (Fig. 6d): 43 of 322 papers with agreed labels mentioned the package without running it, so *U*, the share of mentioning papers that ran it, was 0.87 on average (0.81-0.91). These samples cover only papers that also mention ImageJ/Fiji.

#### Downloads

Each user downloads a package at least once, so downloads per citing paper set an upper limit on *F*× *C* × *P* (Table S13). In 2021-2025, QuPath’s installers were downloaded 131 times per citing paper, inside the proposed range, and Cellpose and StarDist were installed from PyPI about 310 times per citing paper on average. Between 2021 and 2025 the PyPI ratio more than doubled for both. Notebooks, cloud sessions and test pipelines reinstall Python packages every time they run, which may explain the higher and rising PyPI ratios. For ImageJ/Fiji, the discussion reported 500,000 downloads in 2020. Against 12,951 citations that year and our measured *C*, those downloads allow at most 5.3 for *F* × *P* times downloads per user, below the smallest *F* × *P* the discussion proposed, so the download count misses many Fiji users.

#### Updated formula

Counting the papers that mention or cite a package replaces the guessed factor. *N*_*m*_ = *C*× *N*_*c*_ can be counted in full text for each package and year, and applying the measured *C* to all citations extends the count beyond indexed full text. With *U* from reading samples, *N*_*u*_ ≈*F*× *P*× *U* × *N*_*m*_ estimates user-paper occurrences. It counts a person once for each paper, so a count of distinct users also needs the number of papers each user publishes with the package, and applying *U* to other packages, years or cite-only papers assumes that it transfers. The factor this replaces varied more than sixfold between packages and by more than 30% within QuPath and ilastik over five years, whereas *U* stayed within 0.81-0.91 in the three packages we read. Usage logs or surveys are still needed to measure *F* and *P*.

## Code and Data Availability

The main text gives code and data availability for https://github.com/HenriquesLab/mapping-bioimage-software (version 2026.10.31.2). For every validation draw, the repository provides the draw script, the recorded seed and the identifiers of the drawn papers.

In the tables, ZCDL4M/DL4ME abbreviates ZeroCostDL4Mic/DL4MicEverywhere.

**Supplementary Table S1.**
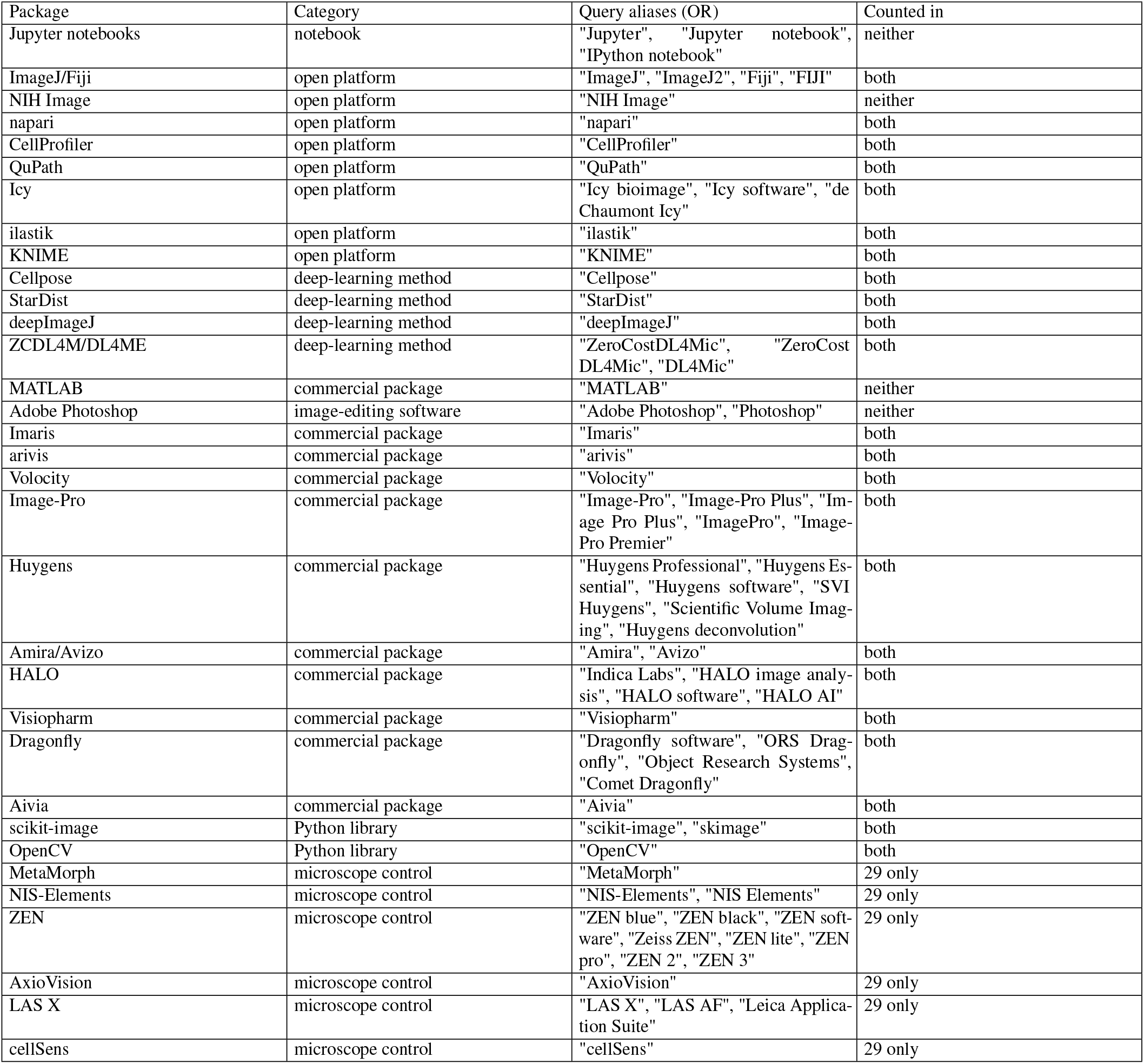
Package registry. Categories and aliases joined by OR in each query; every query also requires microscopy or microscope and indexed full text. “Both” means inclusion in the 23-analysis-package and 29-package totals; “29 only” means microscope-control programs; “neither” means reference, historical-comparison and image-editing entries.

| Package | Category | Query aliases (OR) | Counted in |
|---|---|---|---|
| Jupyter notebooks | notebook | "Jupyter", "Jupyter notebook", "IPython notebook" | neither |
| ImageJ/Fiji | open platform | "ImageJ", "ImageJ2", "Fiji", "FIJI" | both |
| NIH Image | open platform | "NIH Image" | neither |
| napari | open platform | "napari" | both |
| CellProfiler | open platform | "CellProfiler" | both |
| QuPath | open platform | "QuPath" | both |
| Icy | open platform | "Icy bioimage", "Icy software", "de Chaumont Icy" | both |
| ilastik | open platform | "ilastik" | both |
| KNIME | open platform | "KNIME" | both |
| Cellpose | deep-learning method | "Cellpose" | both |
| StarDist | deep-learning method | "StarDist" | both |
| deepImageJ | deep-learning method | "deepImageJ" | both |
| ZCDL4M/DL4ME | deep-learning method | "ZeroCostDL4Mic", "ZeroCost DL4Mic", "DL4Mic" | both |
| MATLAB | commercial package | "MATLAB" | neither |
| Adobe Photoshop | image-editing software | "Adobe Photoshop", "Photoshop" | neither |
| Imaris | commercial package | "Imaris" | both |
| arivis | commercial package | "arivis" | both |
| Volocity | commercial package | "Volocity" | both |
| Image-Pro | commercial package | "Image-Pro", "Image-Pro Plus", "Im-age Pro Plus", "ImagePro", "Image-Pro Premier" | both |
| Huygens | commercial package | "Huygens Professional", "Huygens Es-sential", "Huygens software", "SVI Huygens", "Scientific Volume Imag-ing", "Huygens deconvolution" | both |
| Amira/Avizo | commercial package | "Amira", "Avizo" | both |
| HALO | commercial package | "Indica Labs", "HALO image analy-sis", "HALO software", "HALO AI" | both |
| Visiopharm | commercial package | "Visiopharm" | both |
| Dragonfly | commercial package | "Dragonfly software", "ORS Drag-onfly", "Object Research Systems", "Comet Dragonfly" | both |
| Aivia | commercial package | "Aivia" | both |
| scikit-image | Python library | "scikit-image", "skimage" | both |
| OpenCV | Python library | "OpenCV" | both |
| MetaMorph | microscope control | "MetaMorph" | 29 only |
| NIS-Elements | microscope control | "NIS-Elements", "NIS Elements" | 29 only |
| ZEN | microscope control | "ZEN blue", "ZEN black", "ZEN soft-ware", "Zeiss ZEN", "ZEN lite", "ZEN pro", "ZEN 2", "ZEN 3" | 29 only |
| AxioVision | microscope control | "AxioVision" | 29 only |
| LAS X | microscope control | "LAS X", "LAS AF", "Leica Applica-tion Suite" | 29 only |
| cellSens | microscope control | "cellSens" | 29 only |

Caveats: Jupyter notebooks run other packages. Jupyter and MATLAB are references outside the analysis and field-wide aggregates. MATLAB is used across virtually all quantitative science, so it is excluded from aggregate summaries; its individual count remains reported. Fiji also denotes a country; the microscopy context reduces false matches, although some country matches remain. ImageJ and Fiji count as one ecosystem. Many post-2000 “NIH Image” matches refer to ImageJ, and the modern sample does not estimate precision for the historical program. Icy is a common word, and Huygens also names an optical principle. HALO and ZEN are common words. Dragonfly also names an Andor confocal microscope and an insect. Product phrases and vendor names constrain these queries; Icy, HALO, ZEN and Dragonfly counts are lower bounds. KNIME is a broad analytics platform used beyond imaging; its context clause limits matches to imaging-adjacent literature. Amira is a personal name; one of ten checked matches was an author. napari was released in 2019, so near-zero earlier counts are expected. Every DL4MicEverywhere match also matches a listed alias.

Wildcard additions that are other words come from Table S9. Among 2021-2025 papers citing a package, the citing column gives the share whose indexed text does not mention it, before correction. The last column shows how often the package appears in the full text of 2025 citing papers. Packages without an indexed reference paper have no citation-based miss estimate.

**Sup. Table S2.** Papers mentioning each package. Europe PMC full-text matches in selected years, the 2025 share of all full-text microscopy papers, and 2010-2025 totals from Europe PMC and from OpenAlex titles and abstracts. The last row is the reach denominator.

| Package | 2010 | 2015 | 2019 | 2025 | 2025 share (%) | Europe PMC, 2010–2025 | OpenAlex, 2010–2025 |
|---|---|---|---|---|---|---|---|
| ImageJ/Fiji | 4,289 | 13,221 | 24,248 | 49,255 | 28 | 362,160 | 2,895 |
| MATLAB | 1,450 | 3,561 | 5,396 | 7,331 | 4 | 75,423 | 1,447 |
| ZEN | 59 | 833 | 2,175 | 3,859 | 2 | 28,595 | 51 |
| Adobe Photoshop | 3,218 | 3,677 | 3,434 | 3,220 | 2 | 57,385 | 201 |
| NIS-Elements | 359 | 1,520 | 2,220 | 3,206 | 2 | 31,085 | 112 |
| LAS X | 191 | 904 | 1,625 | 2,761 | 2 | 22,753 | 48 |
| Imaris | 346 | 1,049 | 1,698 | 2,744 | 2 | 24,175 | 270 |
| Image-Pro | 1,046 | 2,127 | 3,026 | 2,641 | 1 | 39,337 | 419 |
| QuPath | 0 | 0 | 65 | 1,625 | 1 | 4,841 | 44 |
| cellSens | 1 | 344 | 704 | 1,352 | 1 | 10,295 | 42 |
| CellProfiler | 41 | 218 | 427 | 1,032 | 1 | 6,911 | 111 |
| Amira/Avizo | 178 | 387 | 574 | 863 | <1 | 8,375 | 157 |
| Cellpose | 0 | 0 | 0 | 778 | <1 | 1,668 | 40 |
| AxioVision | 1,209 | 1,739 | 1,339 | 690 | <1 | 21,551 | 92 |
| MetaMorph | 1,680 | 1,627 | 1,277 | 689 | <1 | 22,269 | 651 |
| Jupyter notebooks | 0 | 6 | 92 | 523 | <1 | 2,291 | 48 |
| OpenCV | 5 | 27 | 103 | 483 | <1 | 2,065 | 64 |
| scikit-image | 0 | 3 | 64 | 443 | <1 | 1,548 | 8 |
| HALO | 0 | 4 | 87 | 442 | <1 | 1,860 | 4 |
| Huygens | 64 | 132 | 316 | 364 | <1 | 3,757 | 7 |
| StarDist | 0 | 0 | 0 | 356 | <1 | 965 | 26 |
| ilastik | 0 | 23 | 101 | 353 | <1 | 1,876 | 49 |
| napari | 0 | 0 | 0 | 278 | <1 | 658 | 38 |
| Volocity | 421 | 763 | 592 | 263 | <1 | 8,966 | 41 |
| Visiopharm | 19 | 73 | 112 | 202 | <1 | 1,641 | 12 |
| arivis | 1 | 3 | 39 | 153 | <1 | 684 | 18 |
| Dragonfly | 0 | 1 | 15 | 119 | <1 | 552 | 17 |
| Aivia | 0 | 0 | 3 | 72 | <1 | 189 | 5 |
| KNIME | 3 | 15 | 47 | 66 | <1 | 609 | 14 |
| Icy | 0 | 12 | 50 | 57 | <1 | 570 | 7 |
| ZCDL4M/DL4ME | 0 | 0 | 0 | 20 | <1 | 116 | 6 |
| deepImageJ | 0 | 0 | 0 | 8 | <1 | 62 | 1 |
| All microscopy papers | 42,806 | 78,337 | 112,489 | 177,358 | 100 | 1,707,893 | – |

**Sup. Table S3.** Deep-learning estimates for 2025. Matches, reads and deep-learning shares; Wilson 95% intervals cover sampling error only for sampled packages. For census rows, Read (n) counts the records in a separately fetched ID list for the same query as Matches, 2025, so the two can differ by a few records.

| Package | Matches, 2025 | Read (n) | DL labels (n) | DL share (95% CI) |
|---|---|---|---|---|
| ZCDL4M/DL4ME | 20 | 19 | 6 | 32% (census) |
| OpenCV | 483 | 484 | 128 | 26% (census) |
| scikit-image | 443 | 440 | 79 | 18% (census) |
| StarDist | 356 | 355 | 53 | 15% (census) |
| napari | 278 | 277 | 39 | 14% (census) |
| Cellpose | 778 | 777 | 106 | 14% (census) |
| Jupyter notebooks | 523 | 523 | 70 | 13% (census) |
| deepImageJ | 8 | 8 | 1 | 13% (census) |
| MATLAB | 7,331 | 200 | 14 | 7% (4-11) |
| ilastik | 353 | 352 | 22 | 6% (census) |
| CellProfiler | 1,032 | 1,032 | 49 | 5% (census) |
| Aivia | 72 | 72 | 3 | 4% (census) |
| QuPath | 1,625 | 1,629 | 62 | 4% (census) |
| Visiopharm | 202 | 203 | 7 | 3% (census) |
| HALO | 442 | 443 | 15 | 3% (census) |
| NIH Image | 34 | 34 | 1 | 3% (census) |
| Amira/Avizo | 863 | 858 | 22 | 3% (census) |
| Icy | 57 | 57 | 1 | 2% (census) |
| Dragonfly | 119 | 119 | 2 | 2% (census) |
| KNIME | 66 | 66 | 1 | 2% (census) |
| NIS-Elements | 3,206 | 200 | 3 | 2% (1-4) |
| Huygens | 364 | 365 | 5 | 1% (census) |
| arivis | 153 | 152 | 2 | 1% (census) |
| LAS X | 2,761 | 200 | 2 | 1% (<1-4) |
| cellSens | 1,352 | 1,356 | 13 | 1% (census) |
| MetaMorph | 689 | 688 | 6 | 1% (census) |
| ImageJ/Fiji | 49,255 | 1,000 | 7 | 1% (<1-1) |
| Imaris | 2,744 | 1,000 | 7 | 1% (<1-1) |
| Adobe Photoshop | 3,220 | 200 | 1 | 1% (<1-3) |
| ZEN | 3,859 | 200 | 1 | 1% (<1-3) |
| AxioVision | 690 | 691 | 3 | <1% (census) |
| Volocity | 263 | 263 | 1 | <1% (census) |
| Image-Pro | 2,641 | 200 | 0 | 0% (0-2) |

**Sup. Table S4.**
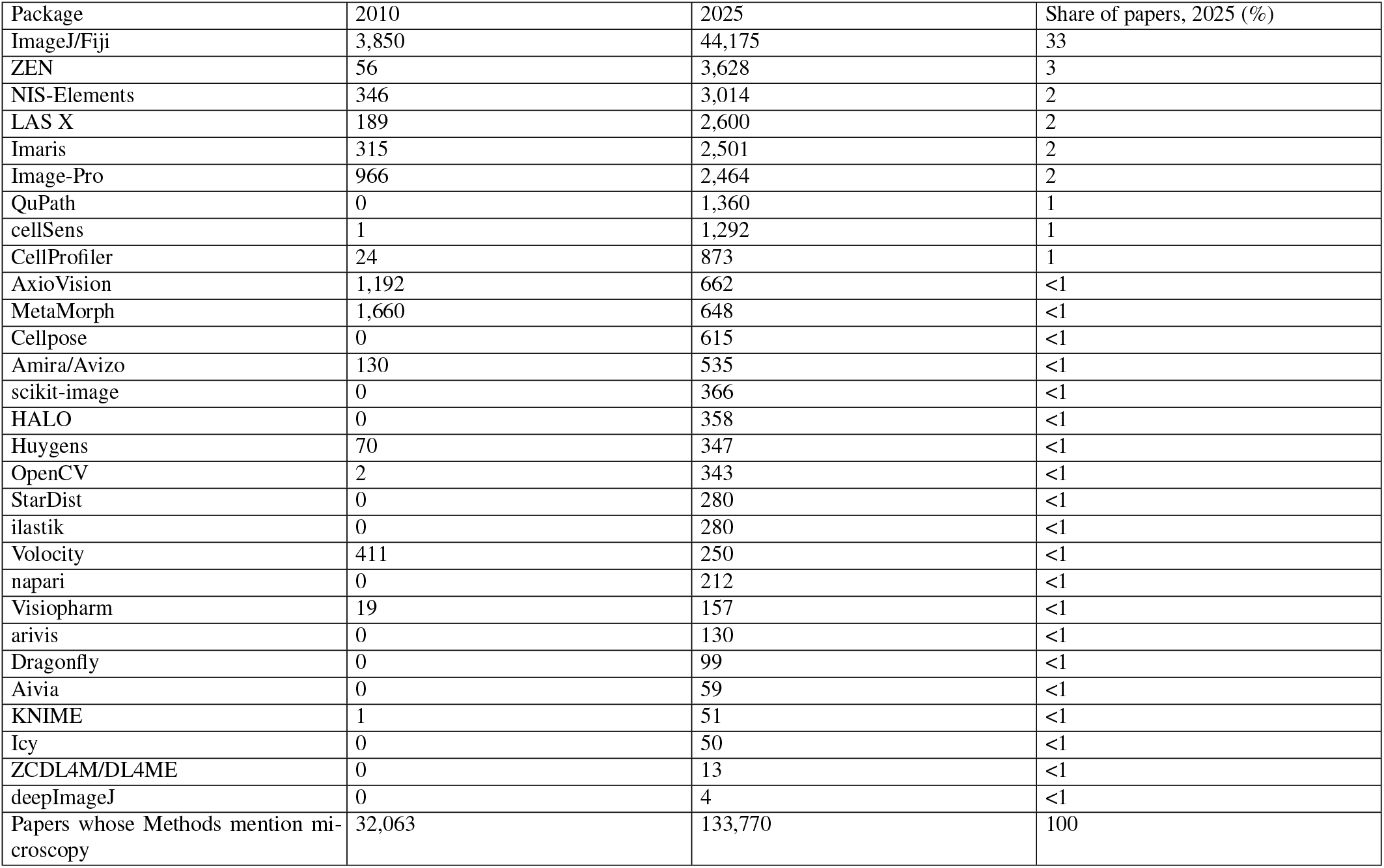
Papers whose Methods mention both the package and microscopy. Counts in 2010 and 2025 and the 2025 share; the last row is the denominator. A matched package may have analysed blots or gels.

| Package | 2010 | 2025 | Share of papers, 2025 (%) |
|---|---|---|---|
| ImageJ/Fiji | 3,850 | 44,175 | 33 |
| ZEN | 56 | 3,628 | 3 |
| NIS-Elements | 346 | 3,014 | 2 |
| LAS X | 189 | 2,600 | 2 |
| Imaris | 315 | 2,501 | 2 |
| Image-Pro | 966 | 2,464 | 2 |
| QuPath | 0 | 1,360 | 1 |
| cellSens | 1 | 1,292 | 1 |
| CellProfiler | 24 | 873 | 1 |
| AxioVision | 1,192 | 662 | <1 |
| MetaMorph | 1,660 | 648 | <1 |
| Cellpose | 0 | 615 | <1 |
| Amira/Avizo | 130 | 535 | <1 |
| scikit-image | 0 | 366 | <1 |
| HALO | 0 | 358 | <1 |
| Huygens | 70 | 347 | <1 |
| OpenCV | 2 | 343 | <1 |
| StarDist | 0 | 280 | <1 |
| ilastik | 0 | 280 | <1 |
| Volocity | 411 | 250 | <1 |
| napari | 0 | 212 | <1 |
| Visiopharm | 19 | 157 | <1 |
| arivis | 0 | 130 | <1 |
| Dragonfly | 0 | 99 | <1 |
| Aivia | 0 | 59 | <1 |
| KNIME | 1 | 51 | <1 |
| Icy | 0 | 50 | <1 |
| ZCDL4M/DL4ME | 0 | 13 | <1 |
| deepImageJ | 0 | 4 | <1 |
| Papers whose Methods mention mi-croscopy | 32,063 | 133,770 | 100 |

**Sup. Table S5.** Validation samples. Each sample has three disjoint replicates; results count items on which both annotators agreed, with the range across replicates in brackets. Workflow samples: run on its own (own), inside Fiji (in Fiji), inside other software (other), combination not stated, or mentioned without use (unused). Per-package name precision is in Table S10.

| Validation check | Sample and replicates | Result |
|---|---|---|
| Aliases, 15 added packages | 3 × 150 | 414 of 440 mention the software (91-96%) |
| Aliases, 15 core packages | 3 × 137-139 | 403 of 412 mention the software (97-99%) |
| Microscope-control use, same matches | 162 agreed | acquisition 68, analysis 90, unclear 4 |
| Lexicon-positive abstracts | 3 × 80 | 215 of 228 deep learning (92-96%) |
| Lexicon-negative abstracts | 3 × 120 | 4 of 342 deep learning (2, 0, 2) |
| Research-area keywords | 3 × 112 | 178 of 297 in the area (55-68%) |
| Cellpose workflow | 3 × 40 | 108 agreed: 68 own, 5 in Fiji, 10 other, 12 not stated, 13 unused |
| StarDist workflow | 3 × 40 | 109 agreed: 19 own, 51 in Fiji, 15 other, 14 not stated, 10 unused |
| ilastik workflow | 3 × 40 | 105 agreed: 67 own, 2 in Fiji, 2 other, 14 not stated, 20 unused |

**Sup. Table S6.** Research-area keywords compared with MEDLINE MeSH headings. Counts give keyword-positive papers with MeSH headings. Agreement gives the share with a mapped MeSH descriptor, and Cohen’s kappa uses both binary labels across all papers with MeSH. Estimates use inverse sampling probabilities, with approximate Wilson confidence intervals based on the conditional Kish effective sample size. The keyword patterns allow a paper to match more than one area. Europe PMC’s MED source does not confirm MEDLINE indexing. Missing headings and differences between keyword vocabulary and indexing limit these associations.

| Area | Papers, n | MeSH agreement (%, 95% CI) | Kappa |
|---|---|---|---|
| Neuroscience | 557 | 61 (57-65) | 0.69 |
| Cancer | 498 | 79 (75-82) | 0.83 |
| Immunology | 685 | 51 (47-55) | 0.56 |
| Microbiology and infection | 407 | 59 (54-64) | 0.66 |
| Development and organoids | 353 | 60 (55-65) | 0.60 |
| Plant biology | 86 | 64 (53-73) | 0.73 |
| Organelles and cytoskeleton | 474 | 57 (52-61) | 0.65 |
| Clinical and pathology | 683 | 12 (9-14) | 0.13 |
| Single-cell and spatial omics | 239 | 30 (24-36) | 0.35 |
| Cardiovascular and renal | 288 | 55 (49-60) | 0.67 |
| Metabolic disease and liver | 204 | 73 (67-79) | 0.81 |
| Electron microscopy | 115 | 43 (34-52) | 0.56 |
| Materials science | 122 | 59 (50-68) | 0.67 |
| Computational methods | 223 | 60 (53-66) | 0.67 |

**Sup. Table S7.** Reporting of image-analysis software in 2025 Methods sections. The share of papers whose sentences mentioning the package give a version, a named function, a parameter, a version with a function or parameter, or none of the three (name only), and the share of articles sharing code. Rates pool three random samples of 200 papers for ImageJ/Fiji, QuPath and CellProfiler and every paper for napari, Cellpose and StarDist, which therefore carry no interval. Brackets give conservative Wilson 95% intervals without finite-population correction.

| Package | Frame (n) | Read (n) | Version (%) | Named function (%) | Parameter (%) | Version and (function or parameter) (%) | Name only (%) | Code shared (%) |
|---|---|---|---|---|---|---|---|---|
| ImageJ/Fiji | 41,322 | 600 | 31 (27-34) | 10 (8-12) | 6 (4-8) | 5 (4-7) | 61 | 14 (11-16) |
| QuPath | 1,251 | 600 | 60 (56-64) | 25 (22-29) | 8 (6-10) | 18 (15-21) | 32 | 27 (23-30) |
| CellProfiler | 764 | 600 | 47 (43-51) | 13 (11-16) | 13 (10-15) | 12 (10-15) | 44 | 39 (35-43) |
| napari | 179 | 179 | 21 | 54 | 6 | 12 | 36 | 78 |
| Cellpose | 517 | 516 | 39 | 40 | 20 | 23 | 38 | 58 |
| StarDist | 244 | 242 | 22 | 22 | 17 | 9 | 52 | 57 |

**Sup. Table S8.** Reporting by computational-methods vocabulary. Rates pool all reporting-audit samples. A paper carries the vocabulary flag when its title or abstract matches the Figure 5 computational-methods pattern. Name only means no version, named function or parameter in the audited sentence window; brackets give name-only counts over the papers in each stratum. The gap compares ImageJ/Fiji with the other packages pooled, as a vocabulary-stratified Mantel-Haenszel risk difference. A paper that mentions several packages counts once for each. In the code repository, XXPROTECTEDBACKTICKXX0XXPROTECTEDBACKTICKXX and XXPROTECTEDBACK-TICKXX1XXPROTECTEDBACKTICKXX record the stratum weights and the papers without title or abstract text.

| Package | With methods vocabulary (%) | Name only, no vocabulary (%, n) | Name only, vocabulary (%, n) |
|---|---|---|---|
| ImageJ/Fiji | 4 | 61 (352/577) | 61 (14/23) |
| QuPath | 9 | 31 (171/545) | 33 (18/55) |
| CellProfiler | 12 | 43 (230/529) | 45 (32/71) |
| napari | 36 | 37 (42/114) | 34 (22/65) |
| Cellpose | 29 | 39 (144/365) | 33 (50/151) |
| StarDist | 26 | 56 (100/179) | 41 (26/63) |

**Sup. Table S9.**
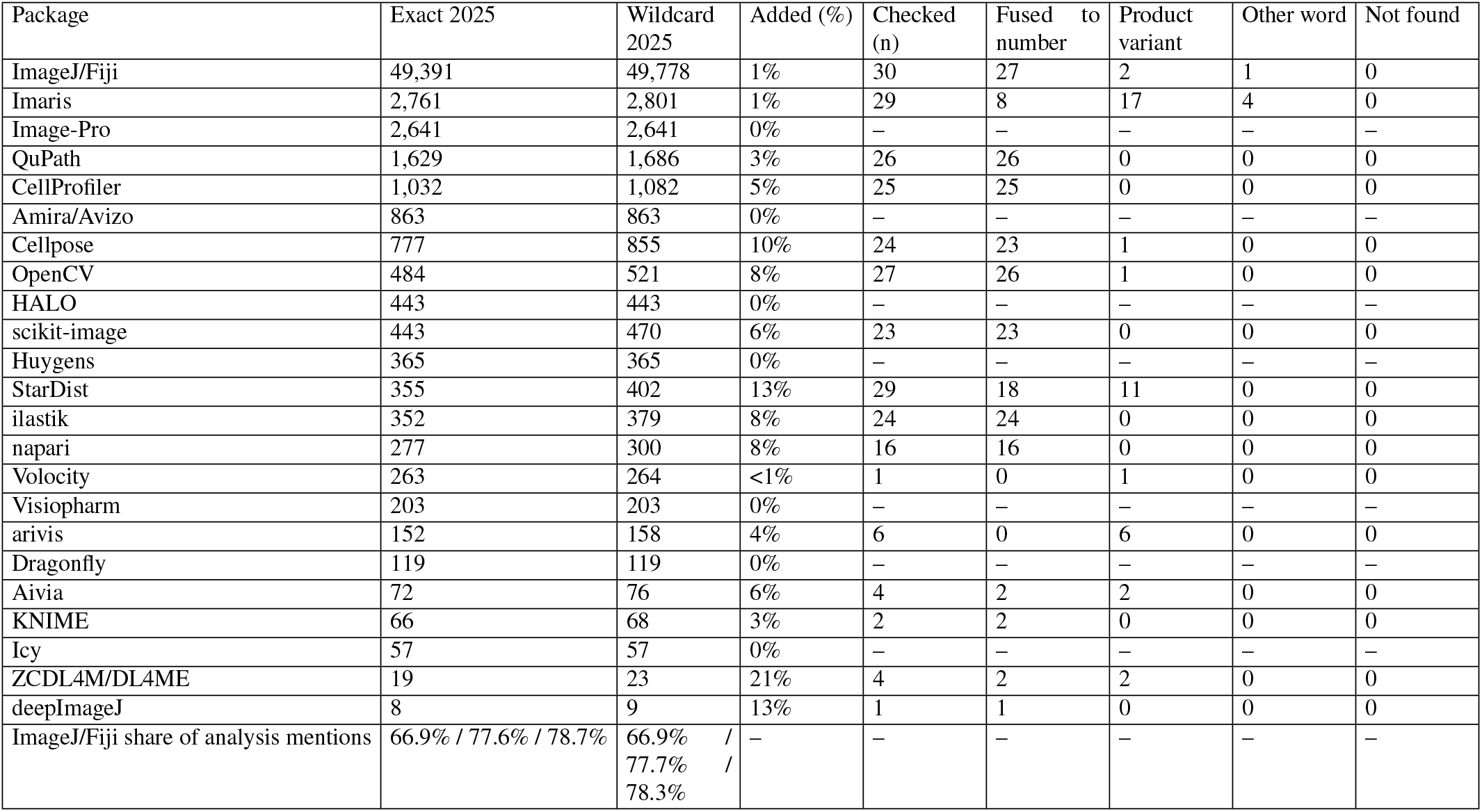
Exact names against trailing wildcards. A trailing wildcard on single-word aliases (QuPath*) catches names the index has joined to a citation number, plus versions and product variants; ordinary words, names and places stay exact (Fiji, Amira, Avizo, MetaMorph, MATLAB, cellSens). Both counts were retrieved on 28 October 2026. For each package, up to 30 of the papers only the wildcard finds were read in full text and the matched word classified. The final row gives the ImageJ/Fiji share of analysis-package matches in 2010, 2020 and 2025, in that order, for exact and wildcard counts, respectively.

| Package | Exact 2025 | Wildcard 2025 | Added (%) | Checked (n) | Fused to number | Product variant | Other word | Not found |
|---|---|---|---|---|---|---|---|---|
| ImageJ/Fiji | 49,391 | 49,778 | 1% | 30 | 27 | 2 | 1 | 0 |
| Imaris | 2,761 | 2,801 | 1% | 29 | 8 | 17 | 4 | 0 |
| Image-Pro | 2,641 | 2,641 | 0% | – | – | – | – | – |
| QuPath | 1,629 | 1,686 | 3% | 26 | 26 | 0 | 0 | 0 |
| CellProfiler | 1,032 | 1,082 | 5% | 25 | 25 | 0 | 0 | 0 |
| Amira/Avizo | 863 | 863 | 0% | – | – | – | – | – |
| Cellpose | 777 | 855 | 10% | 24 | 23 | 1 | 0 | 0 |
| OpenCV | 484 | 521 | 8% | 27 | 26 | 1 | 0 | 0 |
| HALO | 443 | 443 | 0% | – | – | – | – | – |
| scikit-image | 443 | 470 | 6% | 23 | 23 | 0 | 0 | 0 |
| Huygens | 365 | 365 | 0% | – | – | – | – | – |
| StarDist | 355 | 402 | 13% | 29 | 18 | 11 | 0 | 0 |
| ilastik | 352 | 379 | 8% | 24 | 24 | 0 | 0 | 0 |
| napari | 277 | 300 | 8% | 16 | 16 | 0 | 0 | 0 |
| Volocity | 263 | 264 | <1% | 1 | 0 | 1 | 0 | 0 |
| Visiopharm | 203 | 203 | 0% | – | – | – | – | – |
| arivis | 152 | 158 | 4% | 6 | 0 | 6 | 0 | 0 |
| Dragonfly | 119 | 119 | 0% | – | – | – | – | – |
| Aivia | 72 | 76 | 6% | 4 | 2 | 2 | 0 | 0 |
| KNIME | 66 | 68 | 3% | 2 | 2 | 0 | 0 | 0 |
| Icy | 57 | 57 | 0% | – | – | – | – | – |
| ZCDL4M/DL4ME | 19 | 23 | 21% | 4 | 2 | 2 | 0 | 0 |
| deepImageJ | 8 | 9 | 13% | 1 | 1 | 0 | 0 | 0 |
| ImageJ/Fiji share of analysis mentions | 66.9% / 77.6% / 78.7% | 66.9% / 77.7% / 78.3% | – | – | – | – | – | – |

**Sup. Table S10.**
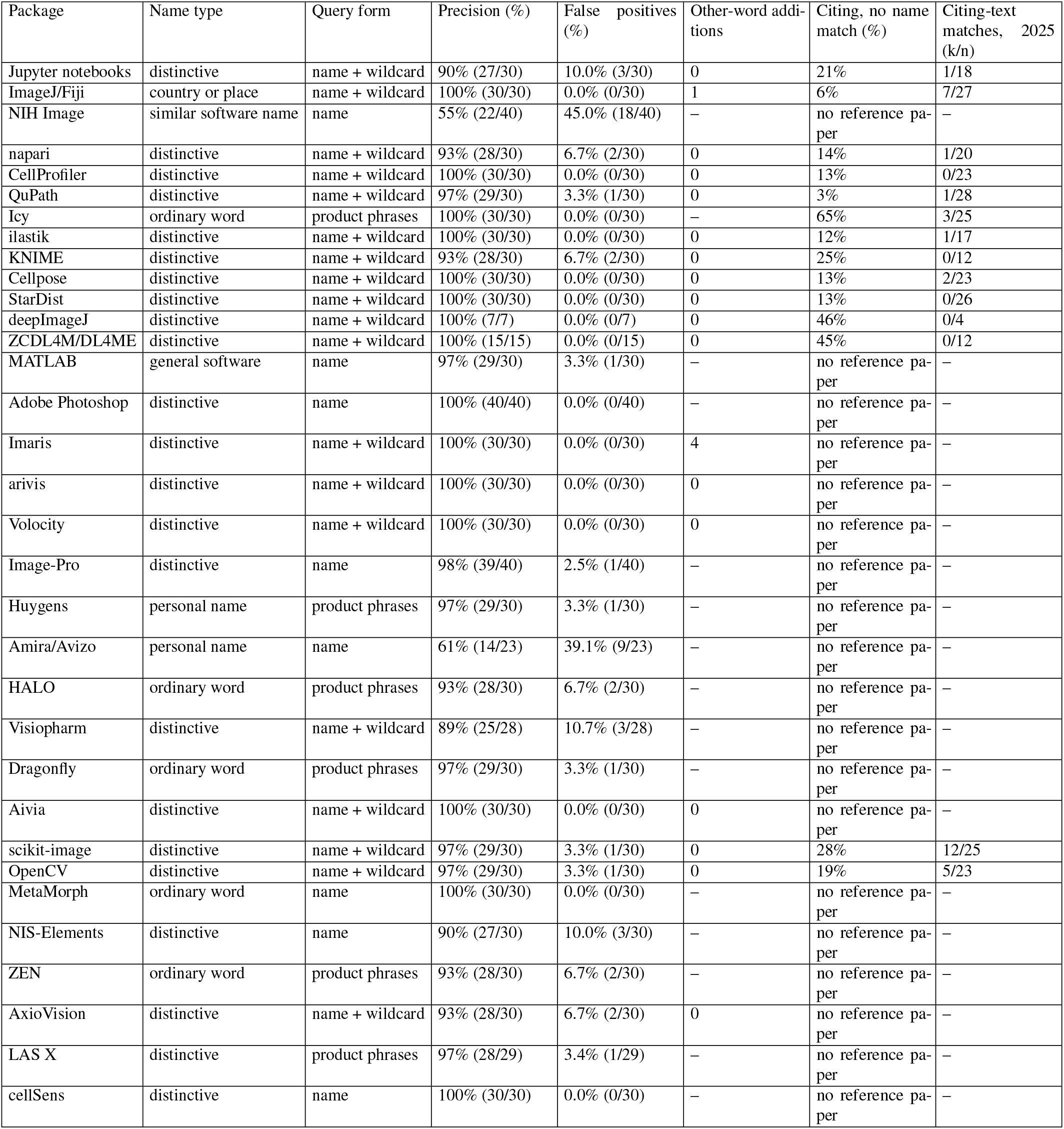
How each package’s query can err. Name type gives the source of ambiguity. For the original registry entries, precision is the share of agreed checked matches judged to mention the software, from three seeded samples of 10 open-access 2025 matches per package. For Image-Pro and Adobe Photoshop, one AI annotator labelled 40 randomly sampled open-access 2025 Methods/full-text snippets per package. NIH Image had separate samples from 2024-2025 and 1995-2003; its query excludes records that mention ImageJ. Image-Pro matched its software in 39 of 40 papers. All 40 Photoshop matches referred to the software; the authors used it for image processing or analysis in 23, figure assembly in 11, illustration in 3, a list of software only in 2 and unclearly in 1. In the 2024-2025 sample, 15 of 40 matches meant the NIH Image program, 19 meant ImageJ, 3 a derivative and 3 were unclear. In the 1995-2003 sample, 22 of 40 meant the program, 3 a derivative and 15 were unclear, and the precision column gives this sample. The repository holds the item-level labels. “Name + wildcard” means the package name with a trailing wildcard in citation analysis.

| Package | Name type | Query form | Precision (%) | False positives (%) | Other-word addi-tions | Citing, no name match (%) | Citing-text matches, 2025 (k/n) |
|---|---|---|---|---|---|---|---|
| Jupyter notebooks | distinctive | name + wildcard | 90% (27/30) | 10.0% (3/30) | 0 | 21% | 1/18 |
| ImageJ/Fiji | country or place | name + wildcard | 100% (30/30) | 0.0% (0/30) | 1 | 6% | 7/27 |
| NIH Image | similar software name | name | 55% (22/40) | 45.0% (18/40) | – | no reference pa-per | – |
| napari | distinctive | name + wildcard | 93% (28/30) | 6.7% (2/30) | 0 | 14% | 1/20 |
| CellProfiler | distinctive | name + wildcard | 100% (30/30) | 0.0% (0/30) | 0 | 13% | 0/23 |
| QuPath | distinctive | name + wildcard | 97% (29/30) | 3.3% (1/30) | 0 | 3% | 1/28 |
| Icy | ordinary word | product phrases | 100% (30/30) | 0.0% (0/30) | – | 65% | 3/25 |
| ilastik | distinctive | name + wildcard | 100% (30/30) | 0.0% (0/30) | 0 | 12% | 1/17 |
| KNIME | distinctive | name + wildcard | 93% (28/30) | 6.7% (2/30) | 0 | 25% | 0/12 |
| Cellpose | distinctive | name + wildcard | 100% (30/30) | 0.0% (0/30) | 0 | 13% | 2/23 |
| StarDist | distinctive | name + wildcard | 100% (30/30) | 0.0% (0/30) | 0 | 13% | 0/26 |
| deepImageJ | distinctive | name + wildcard | 100% (7/7) | 0.0% (0/7) | 0 | 46% | 0/4 |
| ZCDL4M/DL4ME | distinctive | name + wildcard | 100% (15/15) | 0.0% (0/15) | 0 | 45% | 0/12 |
| MATLAB | general software | name | 97% (29/30) | 3.3% (1/30) | – | no reference pa-per | – |
| Adobe Photoshop | distinctive | name | 100% (40/40) | 0.0% (0/40) | – | no reference pa-per | – |
| Imaris | distinctive | name + wildcard | 100% (30/30) | 0.0% (0/30) | 4 | no reference pa-per | – |
| arivis | distinctive | name + wildcard | 100% (30/30) | 0.0% (0/30) | 0 | no reference pa-per | – |
| Volocity | distinctive | name + wildcard | 100% (30/30) | 0.0% (0/30) | 0 | no reference pa-per | – |
| Image-Pro | distinctive | name | 98% (39/40) | 2.5% (1/40) | – | no reference pa-per | – |
| Huygens | personal name | product phrases | 97% (29/30) | 3.3% (1/30) | – | no reference pa-per | – |
| Amira/Avizo | personal name | name | 61% (14/23) | 39.1% (9/23) | – | no reference pa-per | – |
| HALO | ordinary word | product phrases | 93% (28/30) | 6.7% (2/30) | – | no reference pa-per | – |
| Visiopharm | distinctive | name + wildcard | 89% (25/28) | 10.7% (3/28) | – | no reference pa-per | – |
| Dragonfly | ordinary word | product phrases | 97% (29/30) | 3.3% (1/30) | – | no reference pa-per | – |
| Aivia | distinctive | name + wildcard | 100% (30/30) | 0.0% (0/30) | 0 | no reference pa-per | – |
| scikit-image | distinctive | name + wildcard | 97% (29/30) | 3.3% (1/30) | 0 | 28% | 12/25 |
| OpenCV | distinctive | name + wildcard | 97% (29/30) | 3.3% (1/30) | 0 | 19% | 5/23 |
| MetaMorph | ordinary word | name | 100% (30/30) | 0.0% (0/30) | – | no reference pa-per | – |
| NIS-Elements | distinctive | name | 90% (27/30) | 10.0% (3/30) | – | no reference pa-per | – |
| ZEN | ordinary word | product phrases | 93% (28/30) | 6.7% (2/30) | – | no reference pa-per | – |
| AxioVision | distinctive | name + wildcard | 93% (28/30) | 6.7% (2/30) | 0 | no reference pa-per | – |
| LAS X | distinctive | product phrases | 97% (28/29) | 3.4% (1/29) | – | no reference pa-per | – |
| cellSens | distinctive | name | 100% (30/30) | 0.0% (0/30) | – | no reference pa-per | – |

**Sup. Table S11.**
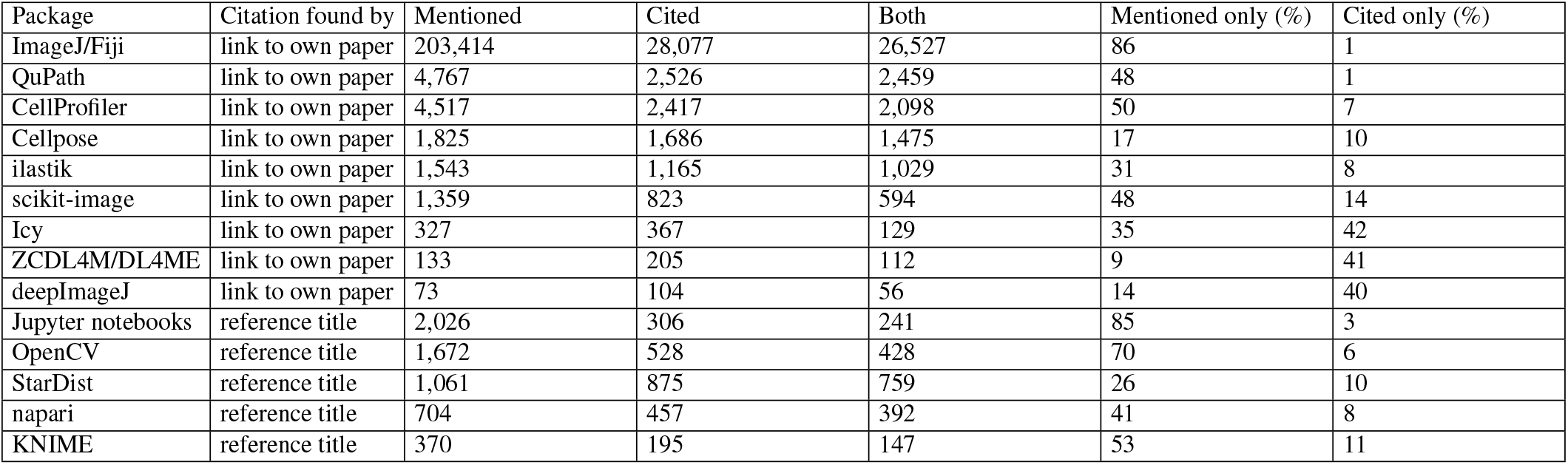
Mentioning and citing each open-source package, 2021-2025. Full-text microscopy papers that mention the package, cite it, or both. For nine packages, a citation is a Europe PMC link to one of the package’s own papers. The papers of napari, StarDist, Jupyter, KNIME and OpenCV are not indexed, so their citations are matches of the package name in reference titles, which also count plugin papers mentioned after a host. Shares are of the papers that mention or cite the package.

**Sup. Table S12.** Mentioning without citing in preprints and journal papers, 2021-2025. For the nine packages with citation links, the share of papers mentioning or citing the package that mention it without citing its papers, among records whose reference list Europe PMC has parsed. Each cell gives the papers mentioning the package and, in brackets, the share mentioning it without citing its papers. The last row gives the parsed share among all full-text microscopy records; a record without a parsed list cannot show a citation link, so including such records would inflate the share for preprints.

| Package | Preprints (n; uncited %) | Journal papers (n; uncited %) |
|---|---|---|
| ImageJ/Fiji | 2,750 (70) | 198,385 (86) |
| QuPath | 167 (56) | 4,451 (46) |
| CellProfiler | 145 (55) | 4,279 (49) |
| Cellpose | 140 (21) | 1,625 (14) |
| ilastik | 79 (32) | 1,406 (28) |
| scikit-image | 88 (50) | 1,219 (46) |
| Icy | 8 (20) | 306 (34) |
| ZCDL4M/DL4ME | 12 (6) | 117 (8) |
| deepImageJ | 3 (0) | 67 (13) |
| Parsed reference lists (%) | 60 | 99 |

**Sup. Table S13.**
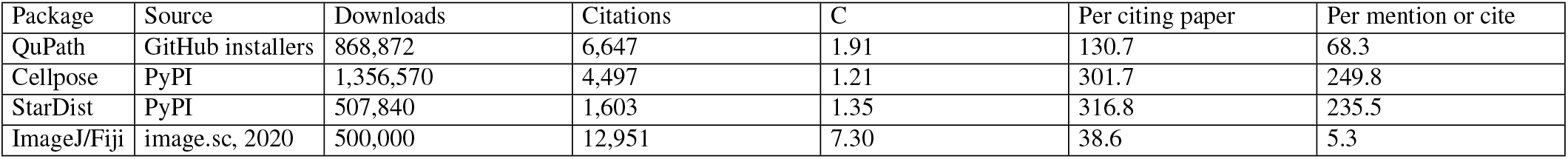
Downloads and citations, 2021-2025. Downloads count PyPI installations by end-user installers (pip, uv, poetry, pdm and pipenv), or downloads of installers attached to GitHub releases published in the period. Citations count all citations of each package’s own papers in OpenAlex, summed over those papers, so a paper citing two of them counts twice. *C* is the measured number of papers mentioning or citing the package for each paper citing it (Table S11). Downloads per citing paper bound the multiplier *F ×C ×P* when each user downloads the package at least once; dividing by *C* bounds *F ×P* times downloads per user. The ImageJ/Fiji row uses the 2020 downloads reported in the image.sc discussion and OpenAlex citations of the three ImageJ and Fiji papers in 2020. Not estimable: CellProfiler and ilastik, whose installers are downloaded from their own websites, which publish no counts; napari, cited mainly through a software record; and scikit-image, OpenCV and Jupyter notebooks, installed mostly as dependencies of other packages.

| Package | Source | Downloads | Citations | C | Per citing paper | Per mention or cite |
|---|---|---|---|---|---|---|
| QuPath | GitHub installers | 868,872 | 6,647 | 1.91 | 130.7 | 68.3 |
| Cellpose | PyPI | 1,356,570 | 4,497 | 1.21 | 301.7 | 249.8 |
| StarDist | PyPI | 507,840 | 1,603 | 1.35 | 316.8 | 235.5 |
| ImageJ/Fiji | image.sc, 2020 | 500,000 | 12,951 | 7.30 | 38.6 | 5.3 |

**Sup. Fig S1.**
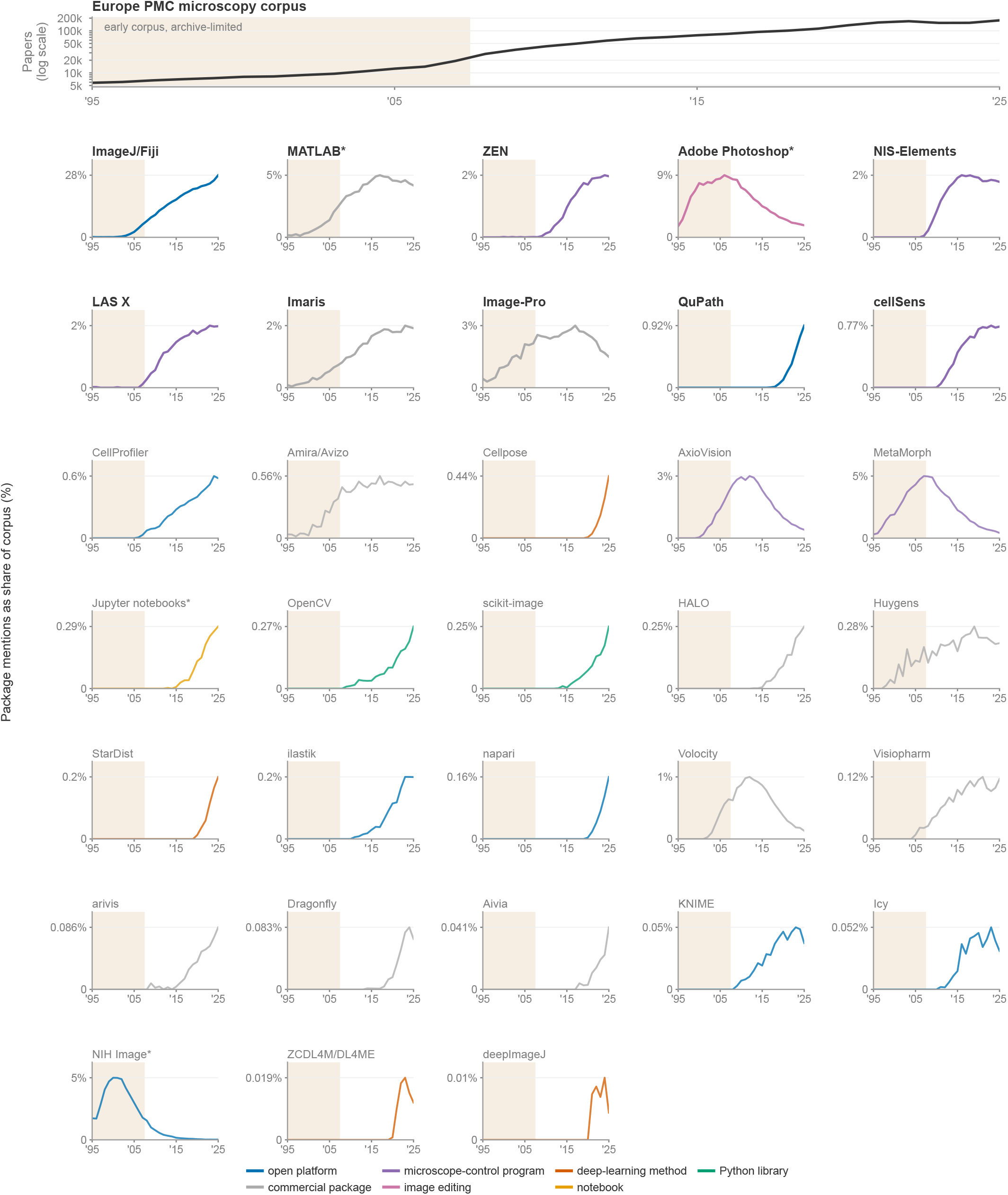
Historical package mentions, 1995-2025. The top panel shows annual Europe PMC microscopy-corpus size on a logarithmic axis. Each lower panel shows the package’s yearly mentions as a share of that corpus; line colour marks its registry category, and each package with mentions appears in its own panel. The ten packages with the most mentions in 2025 have bold labels. Asterisks mark Adobe Photoshop, MATLAB, Jupyter and NIH Image, which enter neither aggregate. After about 2000, many NIH Image mentions refer to ImageJ, so the NIH Image panel overstates the historical program, although its query excludes records that also mention ImageJ. The shaded 1995-2007 zone marks the small corpus dominated by a few journals’ back archives. The corpus and package counts for 1995-2009 and the three new packages’ 2010-2025 counts were queried on 28 October 2026. The 2010-2025 corpus and all other package counts use the 2 October 2026 census snapshot.

**Sup. Fig S2.**
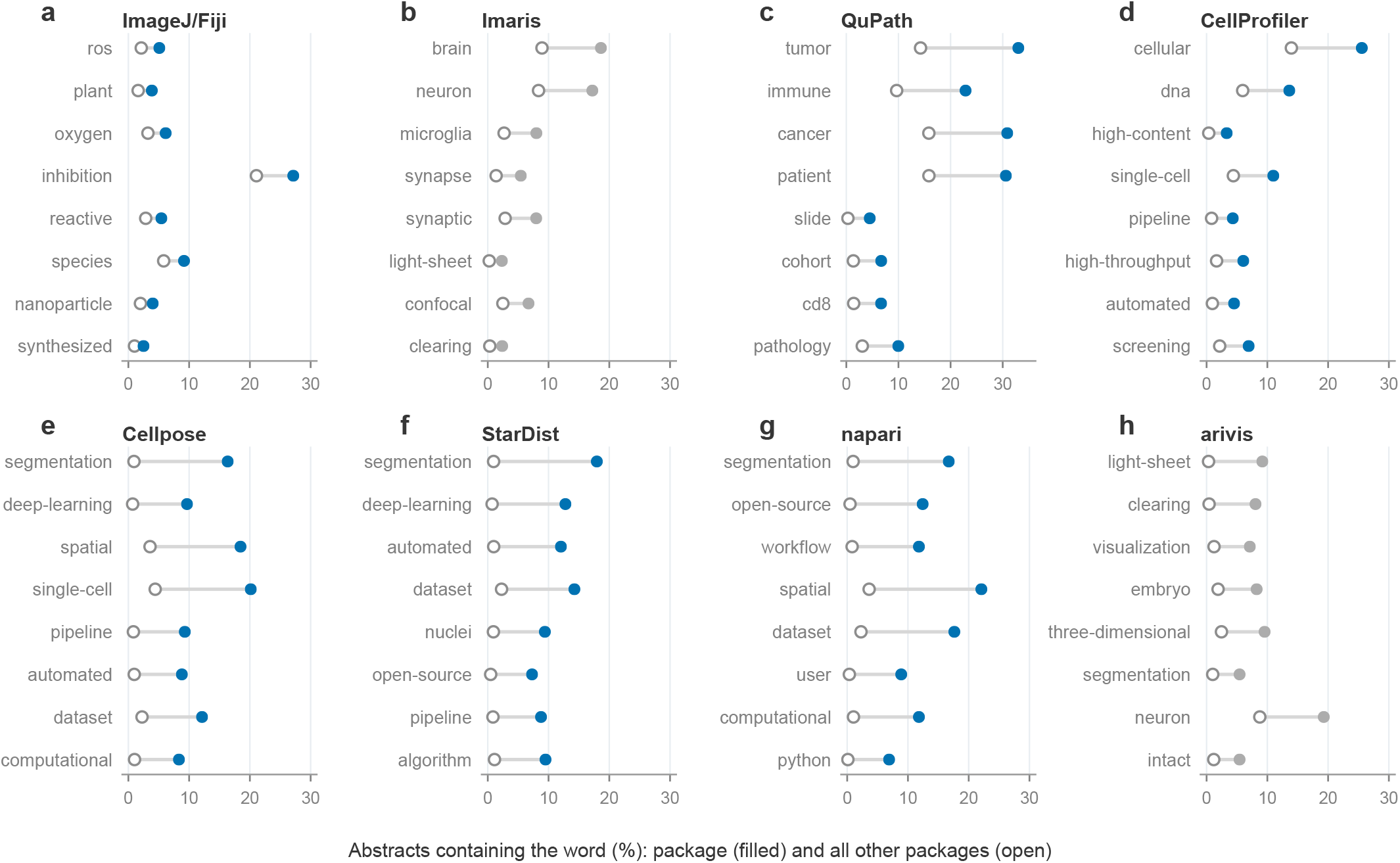
Words over-represented in each package’s papers. For eight packages, the eight words most over-represented in their 2021-2025 abstracts relative to all other analysis packages, among words in at least 2% of the package’s abstracts; word variants and phrases such as light sheet are merged. Filled circles give the share of the package’s abstracts containing the word, open circles the share among other packages. deepImageJ and ZeroCostDL4Mic/DL4MicEverywhere have too few abstracts and are not shown.

